# Concurrent D-loop cleavage by Mus81 and Yen1 yields half-crossover precursors

**DOI:** 10.1101/2023.08.10.552596

**Authors:** Raquel Carreira, F. Javier Aguado, Maria Crugeiras, Marek Sebesta, Lumir Krejci, Miguel G. Blanco

## Abstract

Homologous recombination involves the formation of branched DNA molecules that may interfere with chromosome segregation. To resolve these persistent joint molecules, cells rely on the activation of structure-selective endonucleases (SSEs) during the late stages of the cell cycle. However, the premature activation of SSEs compromises genome integrity, due to untimely processing of replication/recombination intermediates. Here, we employed a biochemical approach to demonstrate that the budding yeast SSEs Mus81 and Yen1 possess the ability to cleave the central recombination intermediate known as the displacement loop or D-loop. Moreover, we confirm that, consistently with previous genetic data, the simultaneous action of Mus81 and Yen1, followed by ligation, is sufficient to recreate the formation of a half-crossover precursor *in vitro.* Our results provide not only mechanistic explanation for the formation of a half-crossover, but also highlight the critical importance for precise regulation of these SSEs to prevent chromosomal rearrangements.

## Introduction

Homologous recombination (HR) is an evolutionary-conserved pathway for the repair of DNA double-stranded breaks (DSB), interstrand crosslinks, and protection and processing of stalled replication forks. It relies on the presence of a homologous sequence as a repair template, primarily during the S and G2 phases of the cell cycle when the sister chromatid is available. However, HR occurring between non-allelic regions or homologous chromosomes can result in detrimental outcomes such as loss of heterozygosity (LOH), translocations, or gross chromosome rearrangements, which have been implicated in human diseases^1,2^.

DSB repair through HR involves several sub-pathways, including synthesis-dependent strand annealing (SDSA), double-strand break repair (DSBR), and break-induced replication (BIR). These pathways involve initial resection of the DNA ends to generate 3′ single-stranded DNA (ssDNA) tails^3^. These ssDNA tails are rapidly bound by replication protein A (RPA), a ssDNA-binding protein, which is subsequently replaced by Rad51 recombinase and facilitated by Rad52 and Rad51 paralogs in budding yeast^4^. The resulting Rad51-ssDNA nucleofilament enables homology search and DNA strand invasion into a donor template. This invasion event creates a joint molecule known as a displacement-loop (D-loop), which represents the central intermediate in HR repair^2,5–7^. In yeast, Rad54 is required for D-loop formation and consequent Rad51 displacement from the heteroduplex DNA^8–11^, allowing DNA polymerase (Pol) δ access to the invading 3′end^12^. To prevent the formation of undesirable crossovers (COs), extended D-loops can be disrupted by helicases such as Sgs1-Top3-Rmi1 (STR)^13^, Srs2^14,15^, or Mph1^16,17^. The extended, displaced strand anneals with the second end of the original DSB within SDSA pathway, leading exclusively to non-crossovers (NCO). Alternatively, if the displaced strand of the extended D-loop reanneals with the second end of the initial DSB, it may lead to the formation of a double Holliday junction (dHJ)^18^, the characteristic intermediate of the DSBR model. The dHJ can be either dissolved by the STR complex, generating exclusively NCOs^19,20^ or cleaved by structure selective-endonucleases (SSEs), such as Slx1-Slx4, Mus81-Mms4 (Mus81), and/or Yen1, generating both NCOs and COs^21,22^. However, where only one end is available for annealing, as in single-ended DSBs at collapsed replication forks or eroded telomeres, the repair proceeds through BIR^23^. This pathway involves bubble-migration-driven DNA synthesis via D-loop branch migration, leading to the conservative inheritance of newly synthesised DNA^24–26^. Due to this particular mode of synthesis, several sources of genome instability were linked to BIR. Unrestrained DNA synthesis may extend to the end of the chromosome, resulting in extensive LOH or non-reciprocal translocations^27,28^. Asynchrony between leading and lagging strand synthesis leads to the accumulation of ssDNA, increasing the risk of mutagenesis^29–31^. Moreover, multiple rounds of strand invasion during BIR may drive complex genome rearrangements if happening within disperse repeat sequences^32–34^. Premature resolution of BIR intermediates might also result in half-crossover (HC) events^35,36^, which involves fusion between recipient and donor molecules and generation of a new, one-ended DSB on the donor chromosome, potentially resulting in additional rounds of HC formation known as half-crossover cascades (HCCs)^30,37^.

Mus81 and Yen1 have been implicated in the generation of complex chromosomal rearrangements during BIR, including HC events and translocations, by cleaving the D-loop intermediate^34^. Previous studies suggested that HC formation in the absence of Polδ is partially Mus81-dependent^36^, and that premature activation of Yen1 can hinder BIR progression and lead to an increase in chromosomal loss (CL) and HC events^29^. Moreover, these SSEs have recently been implicated in the generation of multi-invasion mediated-rearrangements (MIR), a novel source of genetic instability similar to HCCs^38–41^. While some biochemical evidence exists regarding how Mus81 can process a synthetic oligonucleotide-based D-loop substrate^42–44^, little is known about Yen1’s ability to process such synthetic structures. Moreover, it remains unclear if these SSEs can process more physiologically relevant Rad51-mediated D-loops, where the proteins involved in D-loop formation could modulate or interfere with their activity, as evidenced in related biochemical experiments^13,14,31,45–47^.

In this study, we used a biochemical approach to investigate the processing capabilities of Mus81 and Yen1 on various oligonucleotide-based or Rad51-mediated D-loop structures. Our data demonstrate that Yen1 can effectively process all tested D-loop substrates. In contrast, Mus81 fails to cleave D-loops with a ssDNA-overhang. Moreover, by mapping the cleavage sites of these nucleases on synthetic and enzymatically-reconstituted D-loops, we provide evidence supporting their compatibility with the generation of HC and CL events. Importantly, we observed that the simultaneous actions of Mus81 and Yen1 in plasmid-based assays create a double-nicked D-loop structure that can be ligated to form a direct precursor of a half-crossover product. These biochemical results provide mechanistic explanation for the existing genetic evidence, that the combined actions of Yen1 and Mus81 on a D-loop contribute to the generation of complex genome rearrangements in the context of BIR and MIR.

## Results

### Both Mus81 and Yen1 can cleave oligonucleotide-based D-loops

To compare the ability of Yen1 and Mus81 to process synthetic D-loops, we incubated purified proteins with three different oligo-based D-loop structures (D3, D2, D1, Supplementary Table 2). These substrates were 5′-^32^P-end-labelled on one strand, and the reaction products were analysed using native and denaturing PAGE (Fig. 1 and Supplementary Fig. 1). As these three structures have two branching points, the one at the 5′-side of the invading strand will be referred to as the first branching point and the other one as the second branching point. For the synthetic oligonucleotide-based molecule that recapitulates the nascent D-loop intermediate (D3 D-loops), Mus81 exhibited multiple cuts (position 41-47) on the strand complementary to the invading molecule (oligo 1, Fig. 1b-d) as expected from previous studies^42,43,45,48^. The predominant incision occurred 5 nt from the 5′-side of the first branching point (Fig. 1c-d). Yen1 incisions mapping showed that it can cleave all four strands within the D3 D-loop, albeit with different efficiency. The displaced strand (oligo 2) was predominantly cleaved from nt 51 to 53, with the main incision at position 51, which is 1 nt from the 3′-side of the second branching point (Fig. 1c-d). Yen1 could also nick the invading molecule (oligo 3) at 1 to 3 nt from the first branching point. Some nucleolytic activity was detected at the 5′-end of the strand complementary to the invading oligo (oligo 4), explaining the loss of labelling on the native gel (Fig. 1b). Importantly, these activities were absent when the nuclease-dead mutants (ND) were used. Further analysis using a shorter oligo 4 (oligo 4s) confirmed that Yen1’s activity on the 5′-end depends on the proximity of the 5′-phosphate group to the invasion point (Supplementary Fig. 1a-c).

**Figure 1.**
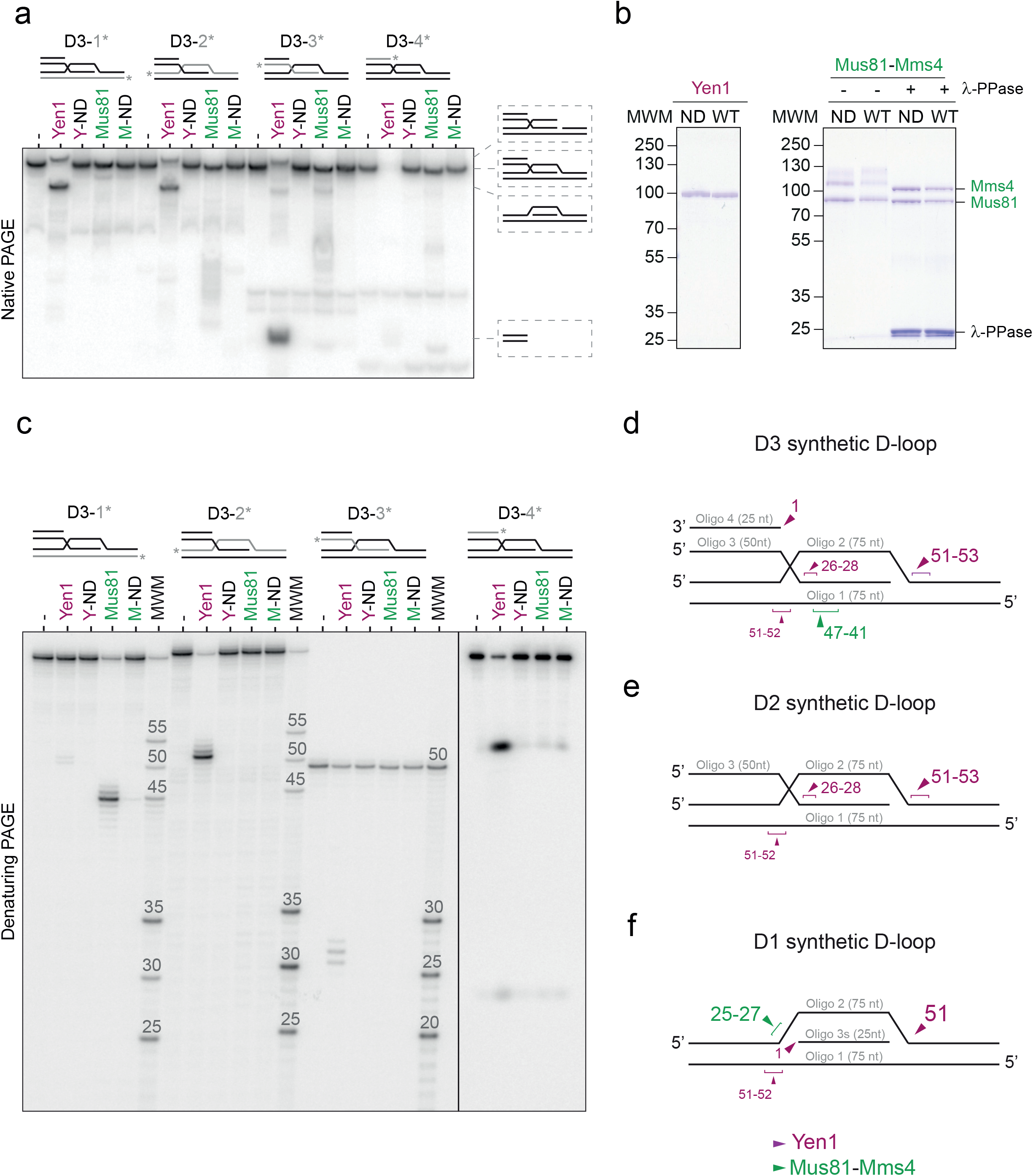
Yen1 and Mus81 cleavage of synthetic D-loops. (**a**) Synthetic D3 D-loops (D3) were ^32^P -labelled at the 5’-end (asterisk) of the indicated strand (grey) and incubated with 10 nM Yen1^WT^ (Yen1), Yen1^ND^ (Y-ND), Mus81^WT^ (Mus81), or Mus81^ND^ (M-ND) for 10 min at 30°C. (-) indicates no enzyme. The reaction products were analysed using 10% native PAGE and phosphorimaged. Schematic representations of the substrate and cleavage products are shown on the right. (**b**) Purified FTH-tagged Yen1^ND^, Yen1^WT^, Mus81-Mms4^ND^ and Mus8-Mms4^WT^ proteins (with or without the lambda-phosphatase (λ-PPase) treatment), were analysed by SDS-PAGE, and stained with Coomassie. Molecular weight markers are indicated in kDa. (**c**) The same reaction products as in (**a**) were analysed using 10% denaturing PAGE. A molecular weight marker (MWM) consisting of a mixture of 5’-^32^P end-labelled oligos of defined length with the same sequence as the labelled oligo was used, therefore MWM bands differ between lanes. D3-4* samples were loaded later on the gel, to allow visualization of small products (**d**) Schematic representation of Yen1 (purple) and Mus81 (green) incision sites on the oligonucleotide-based D3 D-loop. The arrowhead and number size indicate the relative efficiency of the cleavage. (**e**) same as (**d**), but for a D2 D-loop structure. (**f**) Same as (**d**), but for the D1 D-loop structure.

Two additional D-loop structures were generated: the D2 D-loop, characterised by a fully ssDNA 5′-overhang in the invading oligo, which mimics the situation after D-loop bubble migration (Fig. 1e), and the D1 D-loop, a widely used structure for studying recombination intermediates *in vitro*^9,13,16,17,26,49,50^, where the invading strand has no overhangs (Fig. 1f). Yen1 cleaved the D2 D-loop at equivalent positions as D3, while Mus81 was unable to process it (Fig. 1e and Supplementary Fig. 1d-e). In the case of D1 D-loop, both Mus81 and Yen1 cleaved the displaced strand at the branching point. Mus81 predominantly nicked the D1 substrate at position 25 (first branching point), while Yen1 cleaved it 1 nt from the 3’ side of the second branching point (Fig. 1f, Supplementary Fig. 1f-g). Collectively, these results showed that both Yen1 and Mus81 can process oligonucleotide-based D-loop structures according to their respective polarities. Notably, the main incision produced by Mus81 in combination with Yen1’s incision at the end of the displaced strand in D3 structure is compatible with the generation of a HC outcome in the context of BIR repair (Supplementary Fig. 2a). Moreover, Yen1’s cleavage at the invading strand could contribute to the generation of CL events in a BIR scenario (Supplementary Fig. 2b).

### Yen1 incisions on the D3 D-loop are independent

As Yen1 can cut both the invading and the displaced strands within the D3 structure, we aimed to determine if the two predominant Yen1 incisions observed on this structure were co-dependent or could be uncoupled. Time-course analyses of D3 cleavage by Yen1 revealed that processing of the displaced strand occurred faster than that of the invading one (Fig. 2a, c and d). To investigate if the processing of the invading oligo requires prior incision of the displaced strand, we introduced three phosphorothioate linkages between the nucleotides 50-53 of the oligo 2 to inhibit Yen1 cleavage on the displaced strand. This modification resulted in approximately a 50% reduction in Yen1 activity on the displaced strand (Fig. 2b), but only led to a ∼25% decrease in incisions on the invading oligo (Fig. 2c). These findings suggested that the two incisions are not necessarily coupled. To further confirm this observation, we created a D3 D-loop with a nicked displaced strand that mimics Yen1 incision. If cleavage of the invading strand by Yen1 depended on the initial incision, it should be enhanced with this substrate. However, the cleavage kinetics of the invading strand using both nicked and intact D3 D-loops were very similar (Fig. 2e-g, indicating that the two incisions produced by Yen1 on this substrate are independent.

**Figure 2.**
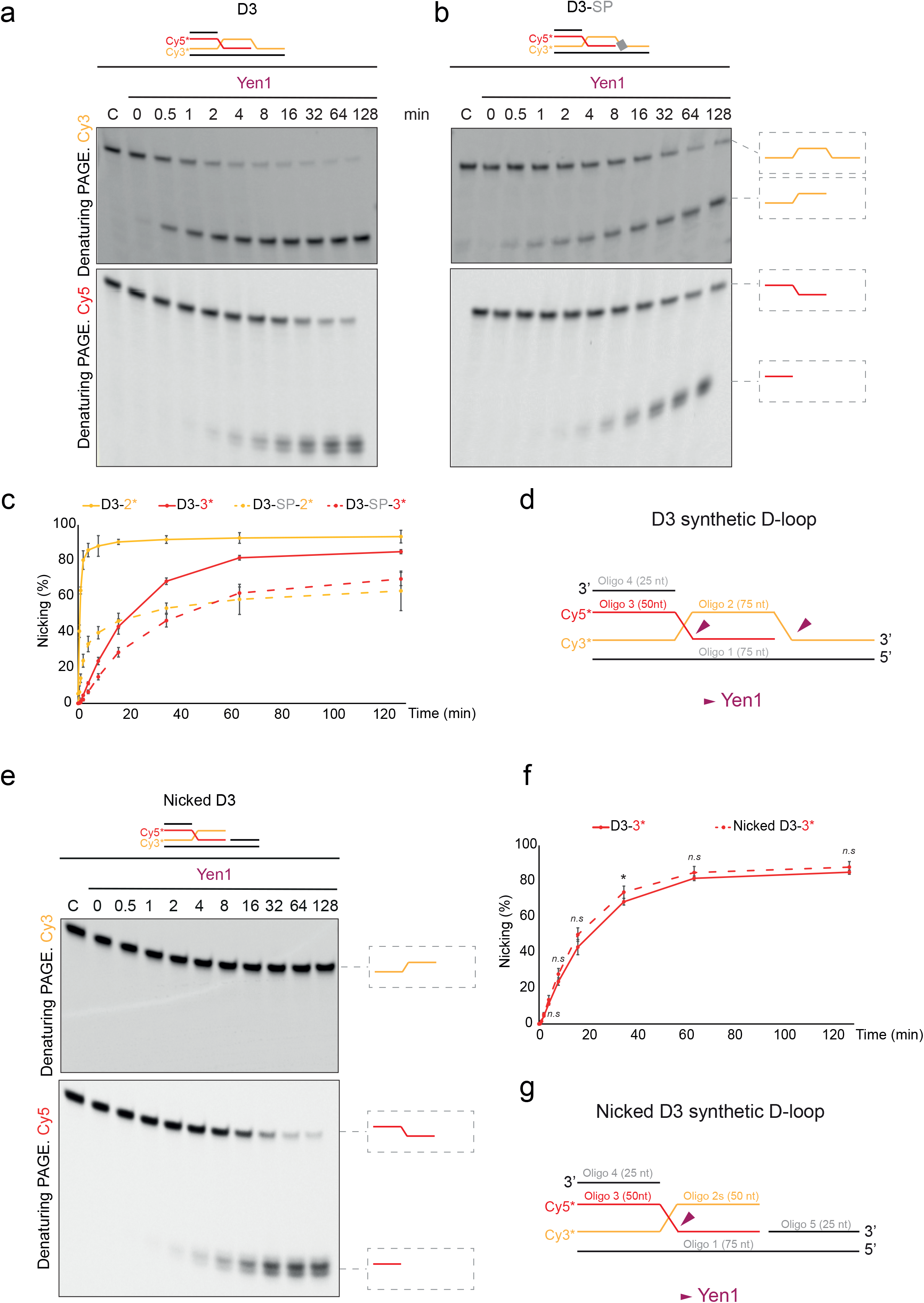
The two incisions produced by Yen1 on a D3 D-loop display different kinetics. (**a**) Time-course analysis of Yen1 cleavage on synthetic D3 D-loop: 10 nM D3 D-loop or (**b**) D3 D-loop with 3 hydrolysis-resistant phosphorothioate linkages (D3-SP, SP linkages in oligo 2 are located between nt 50-51-52-53, depicted by a grey box) were incubated with 20 nM Yen1 at the indicated times at 30°C. The DNA substrates were labelled with Cy3 at the 5′ end of oligo 2 and with Cy5 at the 5′ end of oligo 3. The reaction products were analysed by 10% denaturing PAGE, scanned using a Typhoon FLA9500, and are depicted on the right. Representative gel images are shown. (**c**) Quantification of Yen1-mediated oligonucleotide nicking from denaturing PAGEs shown in (**a**) (solid lines) and (**b**) (dashed lines). Data are represented as mean values ± SD (n = 3). (**d**) Schematic representation of the fluorescent synthetic D3 D-loop structure. Yen1 incision sites are indicated by purple arrowheads. (**e**) Time-course analysis of Yen1 cleavage on nicked D3 D-loop. Reactions were carried out, labelled, and analysed as described in (**a**) and (**b**). (**f**) Quantification of Yen1-mediated oligonucleotide incision of D3 (solid lines) and nicked D3 (dashed lines) D-loop structure. Data are represented as mean values ± SD (n = 3). (**g**) Schematic representation of Yen1 incision (purple arrow) on a nicked D3 D-loop.

### Mus81 and Yen1 can cleave Rad51/Rad54-mediated D-loops

Next, we aimed to address whether Yen1 and Mus81 can also process more physiological D-loops, decorated by the recombination machinery, which may sterically impair nuclease accessibility. For this reason we reconstituted D-loop formation using purified yeast Rad51 and Rad54 (Supplementary Fig. 3a), the negatively supercoiled plasmid (pBSK) as a donor molecule and a fully homologous 6FAM 5′-labelled 90 nt ssDNA (D1) as an invading DNA^49^ (Fig. 3a). After D-loop formation, Yen1 or Mus81 were added, and 15 min later, reactions were stopped, and D-loop processing was analysed by agarose electrophoresis. Both enzymes exhibited dose- and catalysis-dependent destabilisation of the Rad51-mediated D-loop (Fig. 3b), indicating their nucleolytic processing. In Yen1 reactions, a smear dependent on nuclease activity was observed migrating above the free-oligonucleotide band. Considering the exonucleolytic activity detected at the invading oligo in the D1 synthetic D-loop (Fig. 1f), we speculated that this band could result from Yen1 cleaving the 5′-end of the invading strand. This would release 5′-labelled mono-, di- or oligonucleotides with relatively reduced electrophoretic migration, as observed with other fluorochromes^51^. To test this hypothesis, we generated a D1 D-loop using an invading oligonucleotide with three hydrolysis-resistant SP linkages at the 5′-end and incubated it with Yen1 (Supplementary Fig. 3b). In native agarose gels, we observed a slight decrease in the formation of the diffuse band compared to the unmodified D1-Dloop, while denaturing PAGE analysis revealed short cleavage products (5-7 nt) when either substrate was used (Supplementary Fig 3c and d). This suggests that, rather than an exonuclease activity, this smear band might arise from incomplete invasions of the D1 oligonucleotide, leaving short unpaired 5′-ssDNA overhangs that Yen1 could recognise and process as a 5′-flap structure^52^. In addition, to create a more physiological situation, we enzymatically generated two additional D-loops: D2 (with a 5′ ssDNA non-homologous overhang) and D3 (5′ dsDNA non-homologous overhang), which more closely mimic D-loops generated *in vivo*. While Yen1 was able to process both D2 and D3 structures, Mus81 could only act on D3 (Fig. 3c-d), consistent with their processing of the synthetic structures (Fig. 1). In summary, these experiments demonstrate that both Yen1 and Mus81 can process plasmid-based D-loops generated with the proteins involved in their formation *in vivo*.

**Figure 3.**
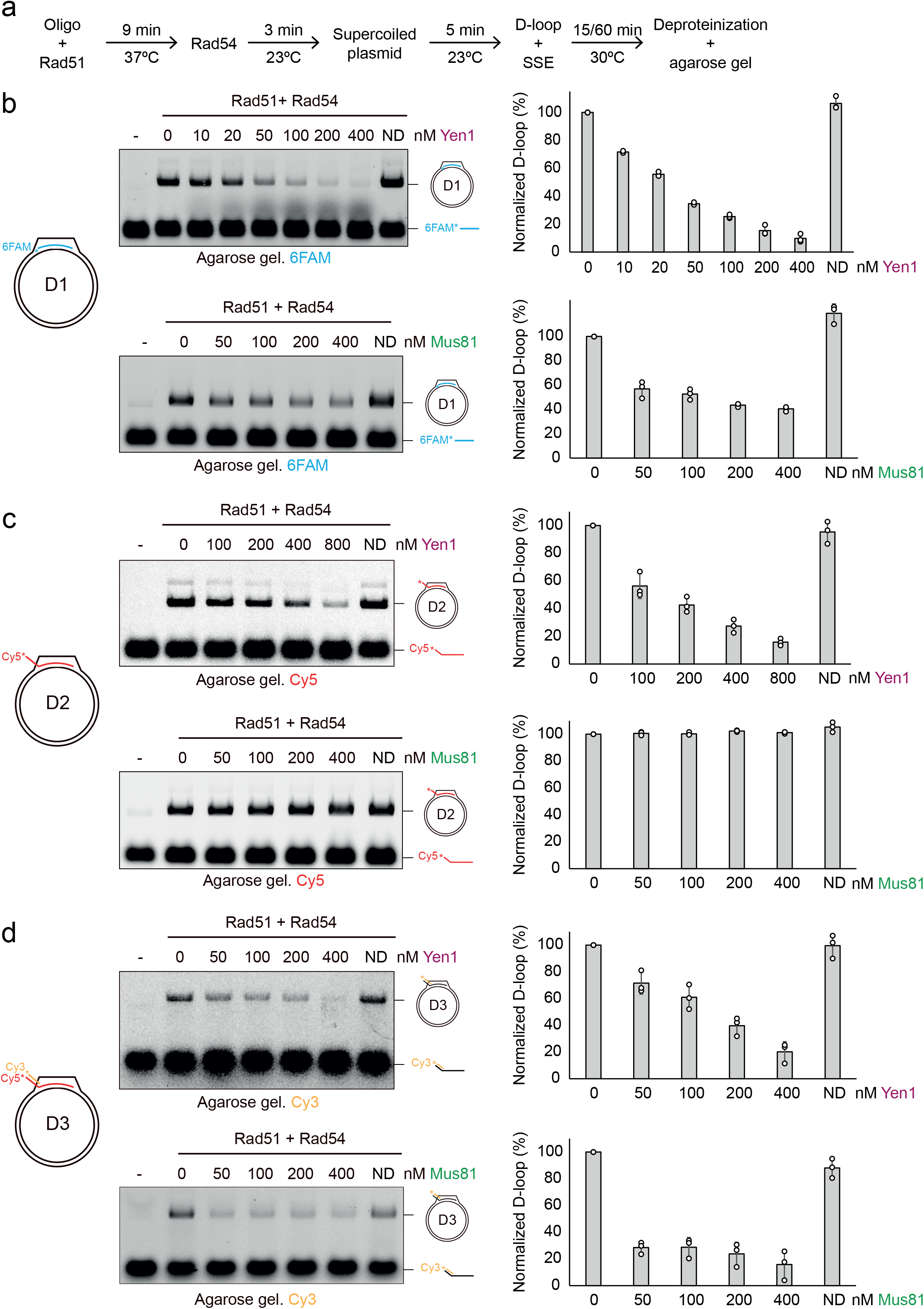
Yen1 and Mus81 process Rad51-mediated D-loops. (**a**) Scheme for Rad51/Rad54-mediated D-loop reaction: End-labelled DNA substrates (40 nM) were incubated with Rad51 (2 µM) at 37°C for 9 min. Rad54 (300 nM) was then added to the reaction and incubated for 3 min at 23°C. Subsequently, supercoiled pBSK (640 ng) was added and incubated for another 5 min before incorporation of SSE. After 15 min incubation with either nuclease at 30°C, the reactions were deproteinised and analysed on agarose gels. (**b**) Representative agarose gel from reactions with enzymatic D1 D-loop treated with Yen1 (top) or Mus81 (bottom). The absence of nuclease is indicated by (-). Nuclease-dead mutant controls (ND) were performed at the highest concentration of the wild-type enzymes. The gels were scanned and quantification of D1 D-loop cleavage by the SSEs is shown on the right. The D-loops were normalised by setting the initial D-loop yield as 100%. The data are plotted as means ± SD (n = 3). White circles represent individual values. A graphical representation of a plasmid-based D1 D-loop is shown on the left. (**c**) Same as (**b**), but using the enzymatic D2 D-loop. (**d**) Same as (**b**) but using the enzymatic D3 D-loop and 60 min for SSE incubation.

### Mus81 and Yen1 cleave Rad51-mediated D-loops coated by RPA

Another protein implicated in D-loop formation in cells is the heterotrimeric ssDNA binding protein (RPA). *In vitro*, RPA stabilises D-loops by binding to the displaced strand and preventing its reannealing to the template strand^53,54^. To check if RPA could influence D-loop cleavage by the SSEs, we generated D1 D-loops in the presence or absence of RPA and used them as substrates for the endonucleases (Fig. 4a). As shown in Fig. 4b, Yen1 cleaves the D1 D-loop with equally efficient regardless of the presence of RPA. In the case of Mus81, rather than preventing cleavage, RPA seems to significantly enhance Mus81 cleavage of the D1 structure (Fig. 4c). Similar experiments with the D3 D-loop confirmed that RPA has no effect on Yen1 activity and mild effect on Mus81 activity on this structure (Fig. 4d-e). To determine if RPA stimulates Mus81 on a simpler substrate, we used a 3′-flap structure with a long ssDNA tail (80 nt) that was pre-incubated for 10 min with increasing concentrations of RPA before adding Mus81 to the reaction. Increasing concentration of RPA inhibited Mus81 activity on this substrate (Supplementary Fig 4a). Importantly, when the same concentration of RPA as used in D-loop reactions was used (600 nM), almost no cleavage was detected. This indicates that RPA not only does not stimulate Mus81 activity on one of its preferred substrates but rather inhibits its nuclease activity, suggesting that observed stimulation of D-loop cleavage might reflect different binding or coordination with RPA. We next addressed if the presence of another single-stranded binding protein on the D1 substrate could stimulate Mus81 activity. To explore this possibility, we performed similar D-loop cleavage assays using the bacterial ortholog of RPA, *E. coli* SSB (Supplementary Fig. 4b). As shown in Supplementary Fig. 4c-d, the presence of SSB did not alter the ability of Mus81 or Yen1 to process D1 D-loops. Therefore, to assess if the stimulatory effect of RPA on Mus81 cleavage might depend on the presence of the Rad51, we generated D1 D-loops using the bacterial ortholog of Rad51, RecA, and subsequently incubated them with SSB or RPA, prior to nuclease incorporation. While both Mus81 and Yen1 were able to process D1 D-loops generated with RecA, no significant differences were observed when SSB or RPA was used (Supplementary Fig. 4e-h). Collectively, these results indicate that neither the presence of RPA nor its bacterial ortholog SSB prevent nuclease activity on the recombination intermediates generated *in vitro*. Furthermore, we have shown that although with very different efficiency, both Yen1 and Mus81 can process RecA-mediated D-loop structures.

**Figure 4.**
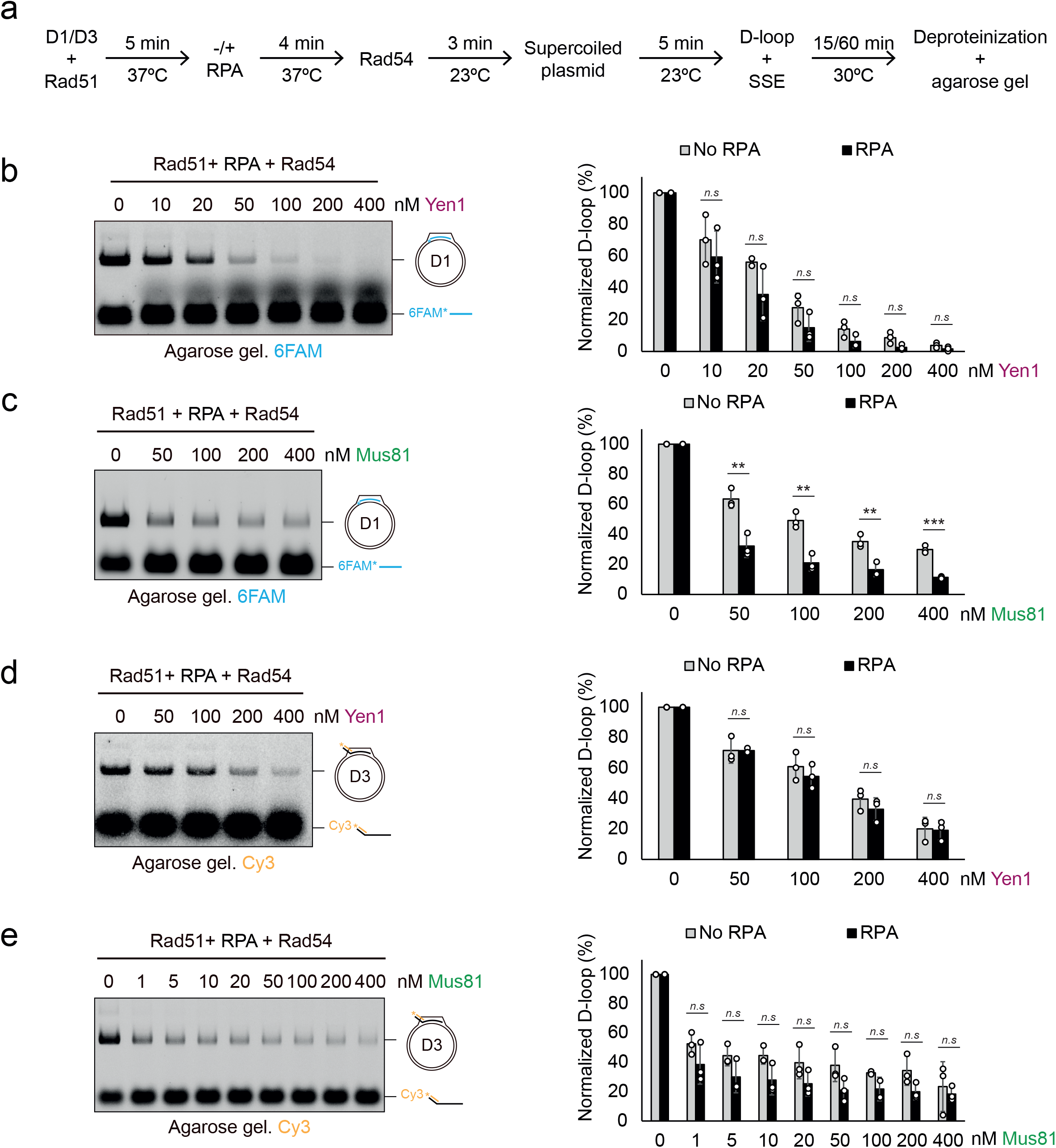
Effect of RPA on D-loop cleavage by SSEs. (**a**) Experimental scheme of Rad51/Rad54-mediated D-loop reactions with RPA. 5′-end-labelled D1 or D3 molecules (40 nM) were incubated with Rad51 (2 µM) at 37°C for 5 min. When appropriate, RPA (600 nM) was added and incubated for 4 min at 37°C. Then, Rad54 (300 nM) was added to the reaction and incubated for 3 min at 23°C, followed by incorporation of supercoiled pBSK (640 ng) and incubation at 23°C for 5 min. The indicated SSE concentrations were added, and reactions were further incubated at 30°C for 15 min for D1 D-loop or 60 min for D3 D-loops. (**b**) Representative agarose gel from reactions with Yen1 and enzymatic D1 D-loop with RPA. Graphs represent the quantification of D-loop cleavage by Yen1. D-loops were normalised by setting the initial D-loop yield as 100% and plotted as means ± SD (n = 3). White circles represent individual values. (**c**) Same as (**b**), but using Mus81. (**d**) and (**e**) Same as (**b**) and (**c**) but for D3 D-loop substrate.

### D-loop proteins impair nuclease cleavage and Rad51 turnover facilitates SSEs activity

Given our previous results, we wanted to investigate if the same proteins required for D-loop formation could modulate the accessibility of the nucleases to the recombination intermediates. To address this, we examined the effect of deproteinisation of Rad51/Rad54-coated D1 structures before Yen1 or Mus81 treatment (Supplementary Fig. 5a). Under identical nuclease concentration, total amount of DNA, and proportion of D-loop formation, both Yen1 and Mus81 displayed faster cleavage kinetics with deproteinised D-loops compared to coated ones (Supplementary Fig. 5b-c). This observation indicates that Rad51 and Rad54 restrict the accessibility of the SSEs to these structures. If the higher efficiency of Yen1 and Mus81 in processing deproteinised D-loops is due to the protective effect of Rad51 and Rad54 proteins, one would predict that generating D-loops with a more stable nucleofilament should reduce accessibility to the nucleases and their catalytic activity. Previous studies have demonstrated that the use of the slowly hydrolysable ATP analogue, ATPγS, leads to a reduction in the efficient turnover of the Rad51-dsDNA complex^55^. We therefore assembled a more stable nucleofilament by incubating Rad51 with ATPγS followed by ATP incorporation upon addition of Rad54 to the reaction (Fig. 5a). As depicted in Fig. 5b, D-loop formation was nearly abolished when only ATPγS was used, whereas incorporation of ATP together with Rad54 restored D-loop formation. Using this strategy, we performed kinetic analyses to examine if the slow turnover of Rad51 at the presynaptic filament could interfere with nuclease cleavage (Fig. 5c). Indeed, stabilisation of the Rad51 nucleofilament using ATPγS strongly reduced the ability of both Yen1 and Mus81 to process D-loop structures (Fig. 5d-e). Taken together, these results indicate that the status of Rad51 nucleoprotein could play a regulatory role against the nucleolytic processing of D-loops by Yen1 and Mus81.

**Figure 5.**
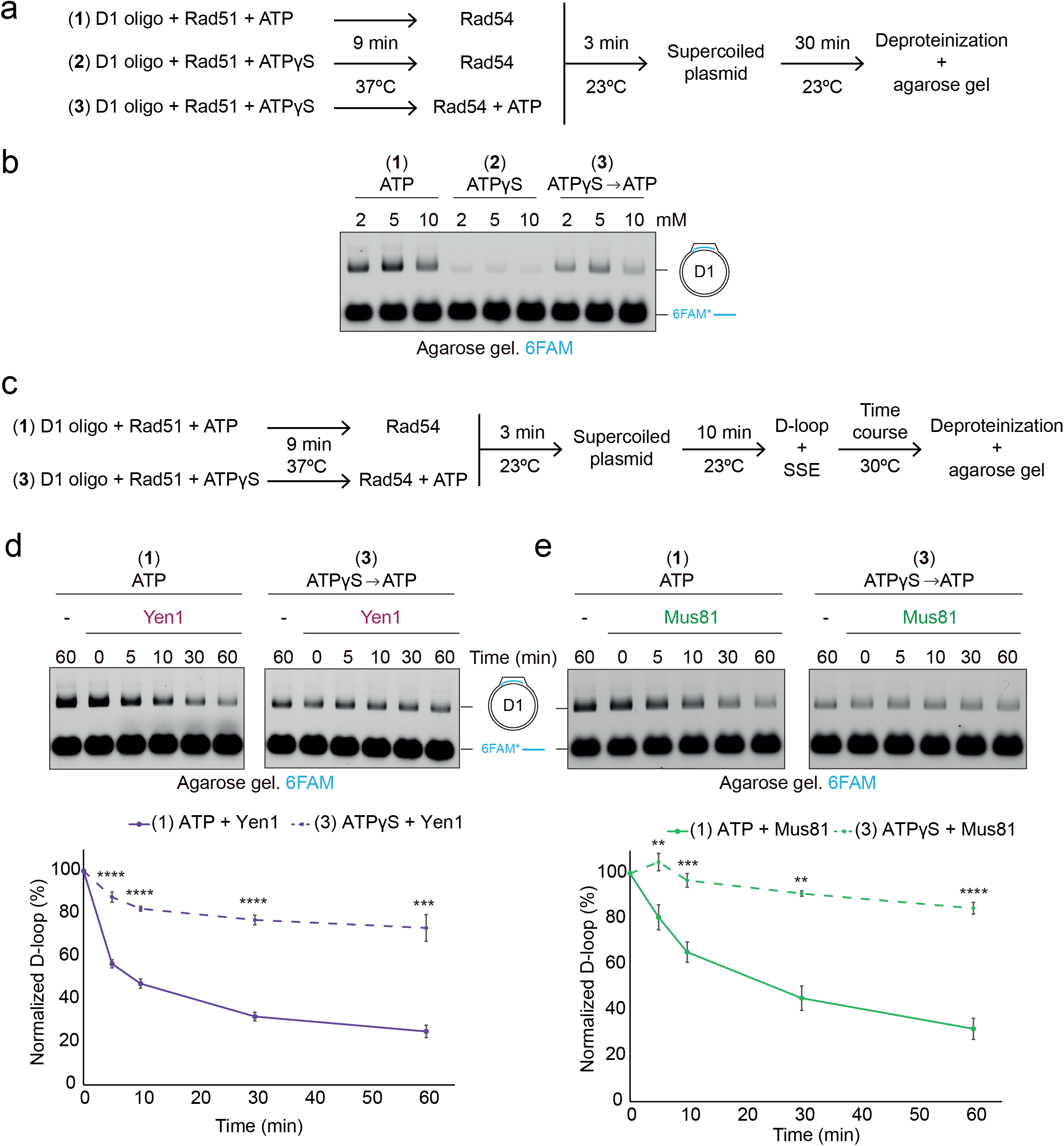
Effect of Rad51 nucleofilament stabilisation on nuclease cleavage. (**a**) Experimental scheme of D1 D-loop reactions with the nucleofilaments generated in the presence of ATP (1), ATPγS (2), or ATPγS followed by ATP incorporation (3). 5′-6FAM-labelled D1 oligo (40 nM) was incubated with Rad51 (2 µM) in presence of either ATP or ATPγS at 37°C for 9 min. Then, Rad54 (300 nM) was added, and in reaction number 3, where ATPγS was used, ATP was incorporated to enable Rad54 activity. Reactions were incubated for 3 min at 23°C. Subsequently, supercoiled pBSK (640 ng) was added and incubated for 30 min at 23°C prior to deproteinization and agarose gel analysis. (**b**) Representative gel of nucleotide effect (2, 5, 10 mM) on D1 D-loop formation in the presence of ATP (1), ATPγS (2), or ATP/ ATPγS (3). Note that when both adenosine nucleotides were used, half of each nucleotide was added to reach of the indicated concentration. (**c**) Experimental scheme of nuclease reactions on D1 D-loops with the nucleofilaments generated in the presence of ATP (1) or ATPγS followed by ATP incorporation (3). Reactions were carried out as stated above. (**d**) Representative agarose gels showing time-course analyses of Yen1 (400 nM) activity on D1 D-loops generated in the presence of ATP (1) (left) or ATPγS/ATP (3) (right). (-) indicates no nuclease. Graph (bottom) represents the quantification of D-loop cleavage by Yen1. D-loops were normalised by setting the initial D-loop yield as 100%. Plotted are means ± SD (n = 3). Solid line: nucleofilament generated in the presence of ATP. Dashed line: nucleofilament generated in the presence of ATPγS. (**e**) Same as (**d**) but using Mus81 (400 nM).

### SSEs incisions on enzymatically made D-loops map at similar positions to those on synthetic D-loops

To address whether the presence of Rad51/Rad54 and RPA not only affects the efficiency of D-loop processing, but also influence the location of incisions produced by these SSEs, we developed a methodology to map their cleavage sites on the plasmid-based D-loops (Fig. 6a). Initially, we focused on the D3 structure and assessed whether the nucleases cleaved the invading strand. After D-loop formation and incubation with Yen1 or Mus81, the reaction products were separated using denaturing PAGE and scanned for Cy5 (invading strand) or Cy3 (complementary oligo in the non-invading duplex region). Yen1 generated a major incision at nt 25 in the 5′-Cy5-labelled invading strand, 5 nt downstream from the beginning of the homology region (Fig. 6b). In addition, several secondary incisions were observed ranging in size from 22 to 40 nt, approximately 20 nt after the start of the homology region (Fig. 6b). This ladder of incisions could be attributed to the dynamics of the D-loop formation process, with Yen1 cleaving the 5′-overhang near the branching point, as observed for the D1 structures (Supplementary Fig. 3d). In contrast, analysis of the cleavage pattern of the 3′-Cy3-labelled oligo revealed no activity (Fig. 6b). Similarly, no activity was detected for Mus81 on neither the invading oligo nor its complementary strand (Fig. 6c), consistent with the results obtained using the synthetic D3 structure (Fig. 1).

**Figure 6.**
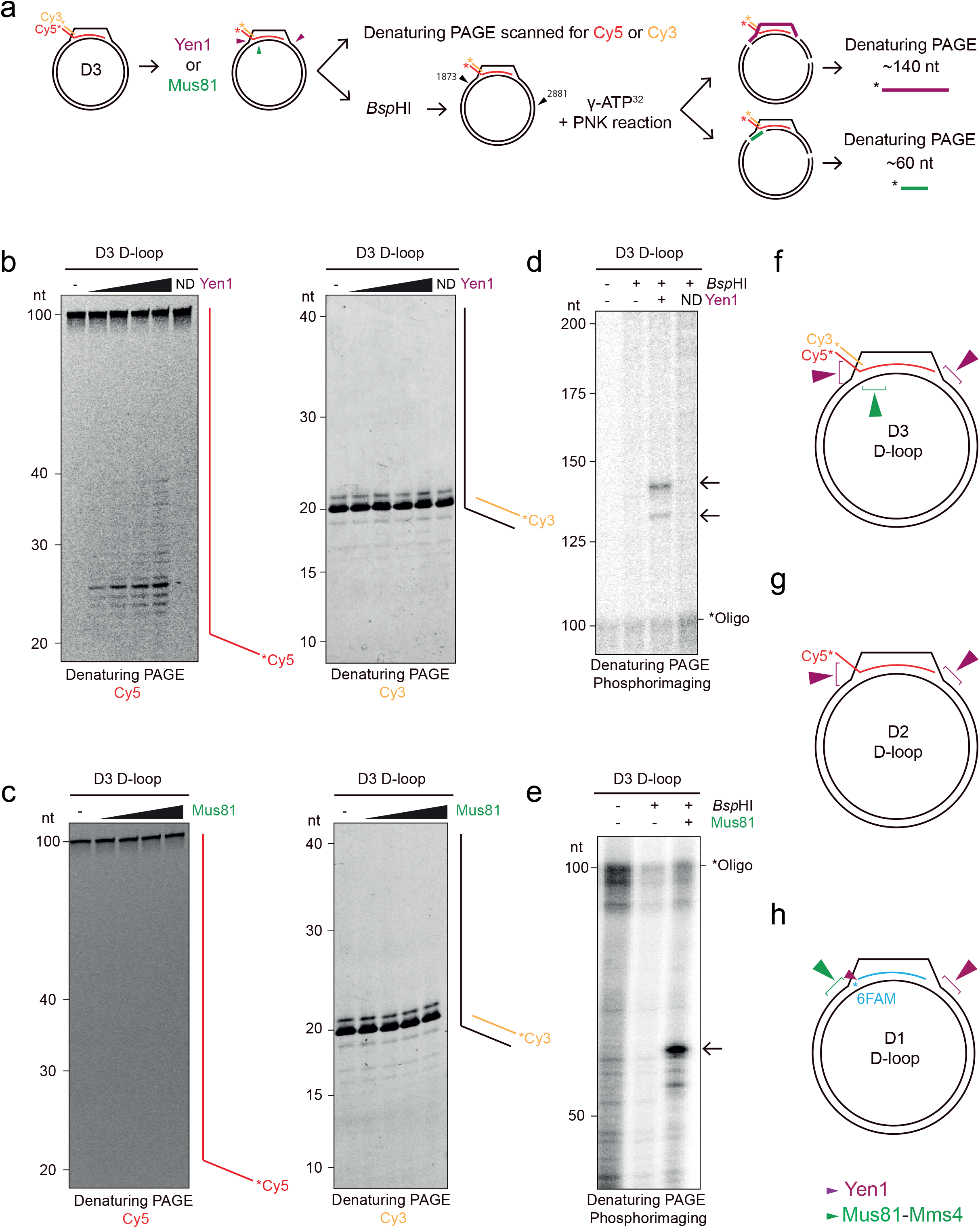
Mapping the D-loop cleavage sites of Yen1 and Mus81. (**a**) Mapping strategy for the D3 D-loop structure. The invading oligo is homologous to positions 1932 to 2012 of pBSK. After D3 D-loop formation, SSE activity on the invading molecule was analysed by fluorescence scanning after denaturing PAGE; SSE activity on the plasmid was analysed after *Bsp*HI treatment, radioactive labelling, and analysis on denaturing PAGE by phosphorimaging. (**b**) To identify cuts in the invading molecule, increasing concentrations of Yen1 (50, 100, 200, 400 nM) or 400 nM Yen1^ND^ (ND) were added to the reaction after D3 D-loop formation and incubated for 1 h at 30°C. (-) indicates no enzyme. Reaction products were analysed by 12% denaturing PAGE and scanned. A mixture of 5’-6FAM-end-labelled oligos of defined length was used as marker. (**c**) Same as (**b**), but using increasing concentrations of Mus81 (50, 100, 200, 400 nM). (**d**) To identify incisions in the plasmid after D3 D-loop formation, 400 nM Yen1 was added to the reaction and incubated for 1 h at 30°C. Then, pBSK was digested with 10 U *Bsp*HI at 37°C for 20 min, followed by treatment with rSAP for another 20 min at 37°C and heat inactivation at 99°C for 5 min. Reaction products were labelled using T4 PNK and ^32^P-γ-ATP and analysed by 6% denaturing PAGE followed by phosphorimaging. A mixture of DNA fragments was radioactively labelled and used as marker. Arrows indicate products of expected sizes. (**e**) Same as (**d**) but using 400 nM Mus81. (**f**) Schematic representation of Yen1 (purple) and Mus81 (green) incisions on the plasmid-based D3 D-loop. (**g**) Same as (**f**) but for D2 D-loop substrate. (**h**) Same as (**f**) but for D1 D-loop substrate.

To determine whether these nucleases were cleaving the plasmid, we treated D3 D-loop reaction products of Yen1 or Mus81 with *Bsp*HI (Fig. 6a, black arrowhead). Given the position of the homology region between the invading oligo and pBSK, an incision by Yen1 at the end of the displaced strand combined with *Bsp*H1 treatment, should release a fragment of approximately 140 nt that could be detected after radioactive labelling and separation by denaturing PAGE (Fig. 6a). Accordingly, only the reactions treated with Yen1 and *Bsp*HI displayed bands near the expected sizes (approx. 140 and 130 nt) (Fig. 6d). Similarly, we confirmed that the combined processing of the D3 substrate by Mus81 and *Bsp*HI released a specific fragment of approximately 60 nt, which corresponds to Mus81 incising the template strand of the plasmid at equivalent positions to the synthetic D3 D-loop (Fig. 6a and e). A limitation of this strategy is its inability to distinguish incisions generated at the expected positions on either strands of the plasmid, as they would release DNA fragments of the same size in combination with *Bsp*HI. Therefore, we complemented these experiments with an alternative mapping approach by utilising the untemplated addition of an adenine nucleotide by *Taq* polymerase at the 3′-end of DNA strands in sequencing reactions (Supplementary Fig. 6a). Consequently, sequencing DNA from SSE-treated D-loop reactions reveal novel A peaks or peaks of increased intensity only in the nicked strand, allowing determination of the strand cleaved by Yen1 and Mus81 with single-nucleotide resolution. As expected, Yen1 cleaved the D3 D-loop at the end of the displaced strand, while Mus81 incised the template strand 4-5 nt upstream of the start of the invasion region, consistent with our observations using the synthetic D3 substrate (Supplementary Fig. 6b). To confirm the specificity of the sequencing results, we conducted similar experiments with a different D3 D-loop structure (D3.2), where the homologous region between the invading strand and pBSK was located at different position (Supplementary Table 2). The sequencing results demonstrated a shift in the A peaks to equivalent positions of the new D3.2 D-loop structure (Supplementary Fig. 6c), thereby validating the mapping approach. The same methodology was applied to determine the incision sites created by Mus81 and Yen1 on D2 and D1 D-loops, respectively (Fig. 6g and h). In the case of the D2 substrate, Yen1 activity resembled its activity on the D3 D-loop, exhibiting multiple cuts on the invading strand and cleaving the displaced strand at the other end of D-loop (Fig. 6g and Supplementary Fig. 7a-d). Further confirmation of the specificity was obtained using a derivate D2 D-loop (D2.2, Supplementary Table 1), in which the 3′end of the invading strand was located at different position within pBSK, resulting in a corresponding shift in the sequencing results (Supplementary Fig. 7e). Regarding the D1 D-loop, Yen1 incisions on the displaced strand were consistent with those observed for D2 and D3. However, Mus81 shifted its activity to the displaced strand, cleaving 5-8 nt upstream of the 5′ end of the homology region (Fig. 6h and Supplementary Fig. 8a-f), consistent with the findings obtained with synthetic substrates. To confirm the mapping of Mus81 incision, an additional D1 oligo (D1.2, Supplementary Table 1) was used, where the 5′-invasion point was again moved to a different position within pBSK (Supplementary Fig. 8g). Altogether, these results suggest that while the Rad51 /Rad54 complex involved in D-loop formation may modulate the accessibility of the nucleases to these structures, its presence does not significantly alter their cleavage specificity compared to the naked oligonucleotide-based D-loops. Furthermore, the newly developed mapping approached enabled precise determination of the incision sites and provided insight into the cleavage preferences of Yen1 and Mus81 during D-loop processing.

### Concurrent cleavage of Mus81 and Yen1 on a plasmid-based D3 D-loop leads to a half-crossover precursor

Our mapping experiments confirmed that Yen1 and Mus81 can introduce incisions on a D3 D-loop that are consistent with the predicted formation of half-crossovers, based on previous genetic studies (Supplementary Fig. 2a)^29,34,36^. Therefore, we next aimed to investigate if the combined activity of these two enzymes could generate such recombination product (Fig. 7a). Surprisingly, when both Mus81 and Yen1 were present simultaneously, a fraction of the D-loop exhibited a shift to a slower migrating band instead of being destabilised (Fig. 7b, arrow). Importantly, the appearance of this new product was dependent on the catalytic activity of both enzymes (Fig. 7c). The reduced migration observed is consistent with the formation of a more open structure containing two nicks, where the invading strand remains paired with pBSK and display a 3′ssDNA flap resulting from Yen1 incision at the end of the displaced strand (Fig. 7d). Based on our mapping, we predicted that this incision would position the 3′-end of the invading strand in close proximity to the newly generated 5′-phosphate end, allowing for theoretical ligation and transfer of the 5’-Cy5 label to the plasmid. To test this hypothesis, we performed subsequent cleavage with *Eco*RV (Fig. 7d), which should generate two linear molecules of approximately 1.2 and 1.7 kb, with the latter being Cy5-labelled (Fig. 7d). Indeed, when we subjected the reaction products from treatment of D3 D-loops with Mus81 and Yen1 to *Eco*RV digestion, we observed the disappearance of the Cy5-labelled band with slower migration and the appearance of a new band of similar intensity of around 1.7 kb (Fig. 7e, left panel, red arrowhead). Ethidium bromide staining of the same gels further revealed the presence of an unlabelled 1.2 kb band (Fig. 7e, right panel, black arrowhead). Importantly, the generation of these products required the combined activity of Yen1, Mus81, and *Eco*RV, but did not require the presence of T4 ligase (Fig.7e, compare lanes 5 and 6 in both panels). To ascertain if the invading strand was indeed ligated to the plasmid, we repeated the experiments and analysed the samples using denaturing PAGE, which should reveal the presence of high molecular weight bands only if the ligation occurred (Fig. 7d and f; 3058 nt expected size: 2958 from pBSK plus 100 nt of D2 oligo). Indeed, the presence of Cy5-labelled DNA was observed in the wells of the gels only when Yen1, Mus81, and T4 ligase were present (Fig. 7f). Further digestion with *Eco*RV enabled the entry of the DNA into the gel, despite its short migration difference (Fig. 7f, compare lanes 5 and 6). To further confirm the production of the expected molecule, we performed the experiments with *Ahd*I digestion, which cleaves closer to the ligation point and thus released a smaller, 129 nt-long Cy5-labelled DNA fragment, which migrates faster in the gel (Fig. 7g, compare lines 8 and 9). These results provide compelling evidence for the ligation of the invading DNA strand to the recipient plasmid, resulting in the formation of a nicked molecule that represents a direct precursor of a half-crossover.

**Figure 7.**
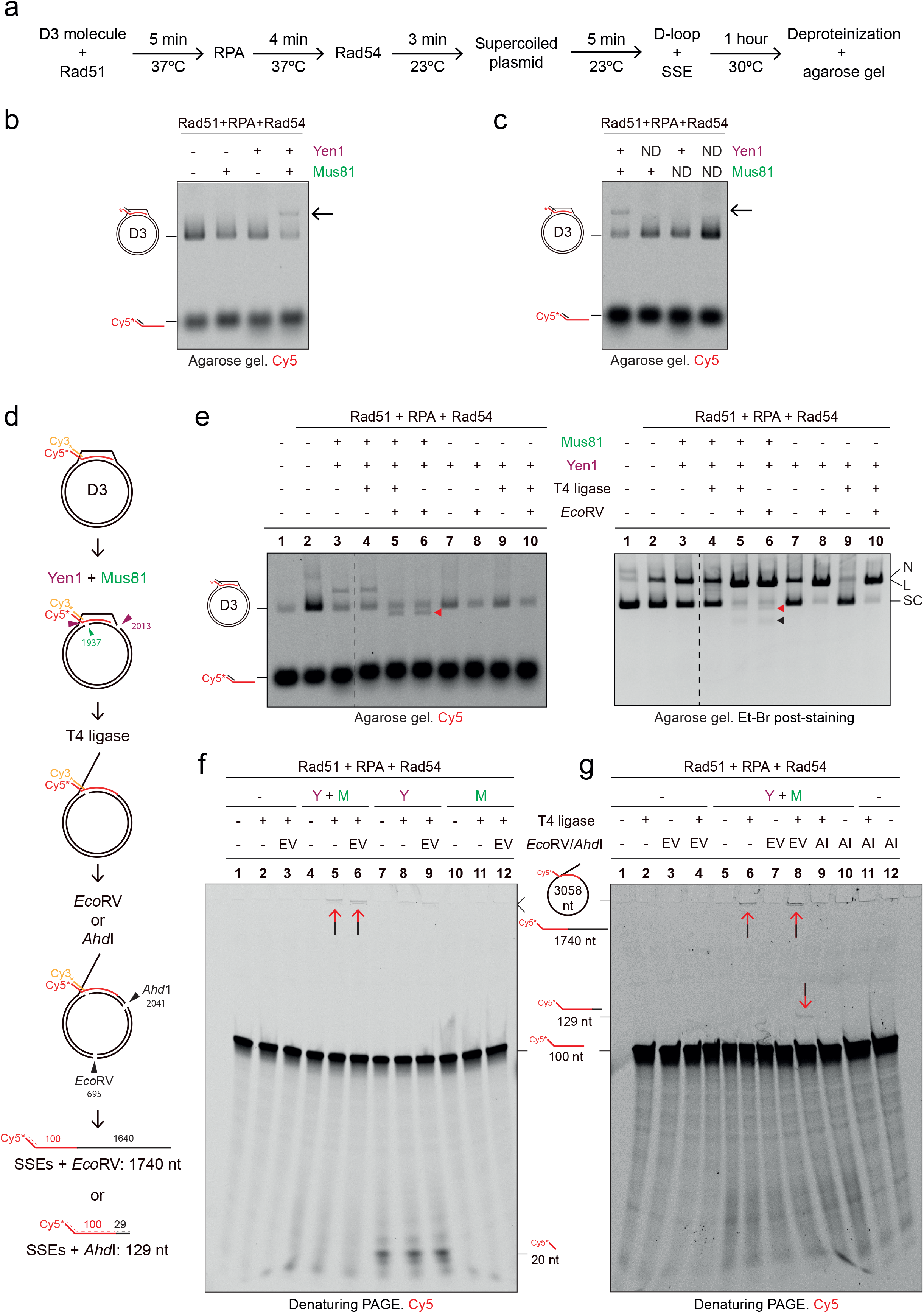
Detection of a half-crossover precursor by concurrent cleavage of D3 D-loop by Mus81 and Yen1. (**a**) Experimental scheme of Rad51-mediated D3 D-loop formation followed by Mus81 and Yen1 resolution. (**b**) Mus81, Yen1, or both were incubated with the D3 D-loop for 1 h at 30°C, resolved on agarose gels and scanned. (-) indicates no nuclease. Arrow indicates product of concurrent cleavage by Yen1 and Mus81. (**c**) Same as (**b**) except nuclease-dead mutants (ND) were used as controls. (**d**) Graphical experimental strategy to detect half-crossover precursors. After D3 D-loop cleavage with Yen1 and Mus81, reaction products were ligated using T4 ligase (1 h, 23°C) and/or cleaved with *Eco*RV or *Ahd*I (1 h, 37°C). After analysis on denaturing PAGE, cleavage by SSEs + *Eco*RV should release a 1740 nt 5′-Cy5-labelled fragment. Cleavage by SSEs + *Ahd*I should release a 129 nt 5′-Cy5-labelled fragment. (**e**) D3 D-loops processing by Mus81 and Yen1 after digestion with *Eco*RV. The red arrowhead indicates the 1716 bp fragment (Cy5 labelled) and black arrowhead indicates the 1242 bp fragment (unlabelled). Cleavage products were analysed on an agarose gel, scanned by a Typhoon FLA9500 (left), and followed by post-staining with EtBr (right). The dashed line indicates gel cropping and removing of irrelevant lanes. Labels: N, Nicked; L, Linear; SC, Supercoiled. (**f**) Reaction products of D3 D-loops processing by Mus81 and Yen1, followed by DNA ligation and cleavage with *Eco*RV were analysed on a 10% denaturing PAGE. The dual coloured arrows indicate the invading strand ligated to the plasmid backbone. Expected products are depicted on the right. Labels: Y, Yen1; M, Mus81; EV, *EcoR*V. (**g**) Same as (**f**) but using *Ahd*I restriction enzyme for product analysis. Labels: Y, Yen1; M, Mus81; EV, *EcoR*V; AI, *Ahd*I.

## Discussion

Genetic data from several groups have implicated Mus81 and Yen1 structure-selective endonucleases (SSE) in the processing of repair intermediates involved in or leading to BIR^26,56–58^. Interestingly, it has also been suggested that the processing of early recombination intermediates, such as D-loops by Mus81 and Yen1 could result in the formation of aberrant repair outcomes, including chromosomal loss and half-crossover events, implying a need for thorough regulation of the SSEs activities^29,34,36,56^. Using a biochemical approach we provide mechanistic insight supporting these observations by comprehensively characterising the activtiy of Mus81 and Yen1 on various D-loop structures.

To mimic different stages of D-loop formation, we examined a nascent D-loop (D3), an extended D-loop (D2), and D-loop with no overhangs (D1), which are widely used models for analysis of *in vitro* D-loop processing. Generally, the cleavage sites detected for Mus81 on the synthetic D3 D-loop (Fig. 1) are in agreement with the previous reports^42–44^, as well as the absence of activity on the D2 D-loop^42^. For the D1 D-loop, incisions happen on the opposite strand (the displaced strand) compared to the D3 D-loop (Fig. 1 and Supplementary Fig. 1), reflecting similarity to a 3’-flap-like structure cleaved by Mus81^43^. For Yen1, we demonstrate for the first time its ability to process all three types of D-loops. We identified an invariant cleavage site on the distal end of the displaced strand in all D-loop strucutures, and observed Yen1’s ability to incise the invading strands (Fig. 1). The locations of these cleavage sites are consistent with the expected 5’ polarity of a Rad2/XPG-family nuclease.

Importantly, the main incision sites remain unchanged when similar D-loop structures are enzymatically reconstituted in a plasmid-based assay (Fig. 3, 6 and Supplementary Fig. 6-8). Nevertheless, D-loop deproteinization prior to nuclease action facilitates the cleavage of both Yen1 and Mus81 (Supplementary Fig. 5), suggesting that Rad51, Rad54, and RPA may influence substrate accessibility to the nucleases, although they do not significantly alter the position of the incisions. Indeed, stabilisation of Rad51 filament in the presence of the non-hydrolysable ATP analogue ATPγS ^59^ (Fig. 5), severely reduced cleavage by Mus81 and Yen1. These experiments point to a possibility that the ATP binding mode of Rad51 and its regulation by Rad51 mediator proteins (*e.g.*, Rad51 paralogs), may play a role in the regulation of Mus81 and Yen1 processing. Additionally, Rad51 has been previously shown to represent a barrier to the catalytic activity of Mus81^46^. In fact, the same study has also shown that Mus81 interacts with Srs2, enabling Mus81 to access its substrate by removing Rad51 from DNA through the “strippase” activity of Srs2 and stimulating directly the Mus81 catalytic activity^46^.

In line with this, RECQ5, a functional human homologs of Srs2, has been found to be required for the removal of RAD51 from late replication intermediates at common fragile sites to facilitate their processing by MUS81-EME1^60^. Similarly, FBH1, another potential human homolog of Srs2, has been shown to cooperate with MUS81 following replication stress^61^. The Srs2 strippase activity is also crucial to prevent the formation of toxic joint molecules during BIR, where long stretches of ssDNA accumulate and can be bound by Rad51, leading to promiscuous invasions events^29^. Importantly, this work indicates that the expression of Yen1^ON^ (a Yen1 mutant refractory to cell cycle-dependent control^52,62^) increases HC and CL events, but only in the presence of Srs2^29^, suggesting that D-loop-bound Rad51 could prevent their processing by Yen1. These findings collectively highlight the essential role of Rad51 and its accessory proteins in processing of joint molecules and the prevention of undesirable recombination outcomes.

Finally, our study provides biochemical evidence supporting the genetic observations indicating that concurrent actions of Mus81 and Yen1 on D-loop structures may facilitate chromosomal rearrangements that occur in the context of BIR, like HCs^29,35,36^. The specific locations of the incisions generated by Mus81 on the non-displaced strand and Yen1 on the displaced strand of the donor DNA are compatible with the generation of HC products if the invading molecule is ligated to the donor through the nicks created by these SSEs (Fig. 8 and Supplementary Fig. 2a). Indeed, we demonstrate that the addition of a ligase in our reconstituted D-loop reactions, following Mus81 and Yen1 treatment allows ligation of the invading strand to the donor molecule (Fig. 7). However, since Mus81 incisions occur 4-5 nt from the first branching point (Supplementary Fig. 6), the resulting small ssDNA gap would prevent ligation. Congruently, we were unable to detect the second ligation event between the short strand of the invading molecule to the pBSK donor (data not shown). Furthermore, we noticed that when Mus81 and Yen1 are incorporated simultaneously, Yen1 is no longer able to cleave the invading strand (Fig. 7f). This is most likely due to Mus81 incision allowing rotation of the D-loop into a RF-like structure where the invasion point is no longer a branched structure that can be targeted by Yen1. Since cleavage of the invading strand by Yen1 most likely leads to CL events (Fig. 8 and Supplementary Fig. 2b), it would be interesting to address in the future if the absence of Mus81 increases CL events in a BIR system like that described in Elango et al. (2017) when Yen1^ON^ is expressed.

**Figure 8.**
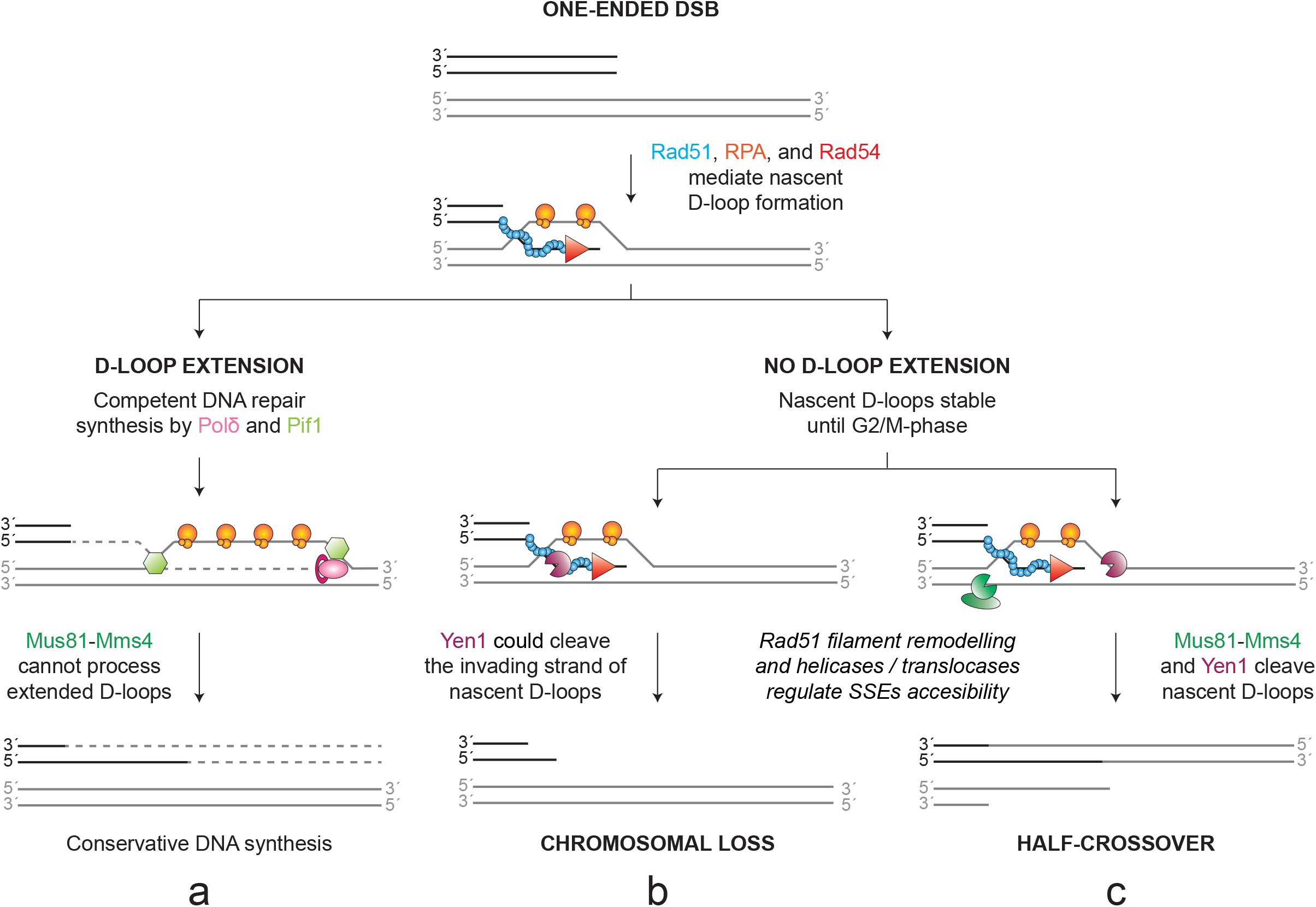
Model for the generation of chromosomal loss and half-crossover events due to Mus81-Mms4 and Yen1 activity on BIR intermediates. Protein names and cartoons are colour-coded. (**a**) After nascent D-loops are formed, de novo DNA synthesis and bubble migration render the D-loop insensitive to Mus81-Mms4 action, allowing their extension until they reach a convergent replication fork or the end of the chromosome (**b**) Persistent nascent D-loop structures that are not extended may become substrates for the SSEs later in the cell cycle or upon premature SSE activation, leading to nascent D-loop cleavage. In this scenario, Yen1 nicking activity at the invading strand may result in a chromosomal loss event. SSE accessibility to this substrate might be regulated by Rad51 filament remodelling, by Rad51 paralogs or any other factor that affects Rad51 ATP hydrolysis. (**c**) As in (**b**), but depicting the combined actions of Mus81-Mms4 and Yen1 leading to a half-crossover event. In addition to Rad51 filament remodelling, several helicases/translocases that have been proposed to modulate Mus81-Mms4 activity could influence the frequency of this outcome (*e.g*., Rad54, Srs2, see discussion section).

Alternatively, using synthetic substrates, we have determined that Yen1 incisions at the equivalent D3 D-loop structure are independent, with the clevage on the displaced strand occuring much faster than the clevage on the invading oligo (Fig. 2). HCs are described to occur when the initiation of BIR through D-loop formation has been initiated, but DNA synthesis is compromised due to defects in Polδ^35–37^ or Pif1^24,26,50,63^, which are both required for extensive DNA synthesis. It has been proposed that the failure to initiate DNA synthesis in these mutants could lead to the resolution of a single HJ generated after strand invasion, and that Mus81 has been suggested to play a role in the generation of these genetic products^36^. In addition, there is a long delay between strand invasion and the initiation of DNA synthesis within the D-loop^64^, which might be due to the activation of a recombination execution checkpoint in the absence of a second end at the DSB^65,66^. In this context, it is plausible that nascent D-loops formed in S or early G2 phases may not be extended until later stages of the cell cycle, when Mus81 and Yen1 become activated^67^. If DNA synthesis within the D-loops begins, the migration of the bubble would separate the invasion point from the complementary strand of the invading molecule^24^, leading to a structure similar to the D2 D-loop. In this configuration, Mus81 would no longer be able to cleave, allowing error-prone BIR synthesis to proceed (Fig. 1, 3 and 8). Moreover, it has been demonstrated that premature SSE activation and entry to mitosis, due to a compromised checkpoint response in cells undergoing BIR, also stimulates HC formation^37^.

Collectively, our findings provide a biochemical framework for the genetic observations implicating Mus81 and Yen1 in the generation of chromosomal loss and half-crossovers events. This study enhances our understanding of the complex interplay between DNA repair pathways and underscores the importance of maintaining genome integrity in safeguarding against genomic instability and disease development.

## Methods

### Recombinant proteins

Yen1 and the catalytically inactive mutant Yen1^ND^ (E193A, E195A) were purified as previously described^68^. Mus81-Mms4 and the catalytically inactive mutant Mus81-Mms4^ND^ (D414A, D415A) were purified as described elsewhere^69^. Rad51, RPA, and Rad54 were purified according to described procedures^49^. RecA was obtained from New England Biolabs (#M0249S) and SSB from Thermo Fisher Scientific (#70032Z500G). Protein concentrations were determined using the Bradford assay (Bio-Rad) and densitometry of Coomassie-stained PAGE gels, using bovine serum albumin (BSA) as a standard. All protein batches were tested for exonuclease, endonuclease, and protease contaminant activities, and were analysed by SDS-PAGE followed by Coomassie staining (Fig. 1b and Supplementary Fig. 3a).

### Oligonucleotide purification and annealing into DNA substrates

The oligonucleotides used in this study (Supplementary Table 1) were obtained from Merk, subjected to PAGE purification, and annealed as detailed previously^68^. Shortly, labelled and unlabelled oligonucleotides were mixed at a 1:3 ratio, boiled in a water bath, and cooled down to room temperature overnight. Fully annealed substrates were purified from 10% native PAGE gels in 1xTris-Borate-EDTA (1xTBE) buffer (90 mM Boric acid, 90 mM Tris-base, 2 mM EDTA). For radioactive substrates, oligonucleotides were 5’-end-labelled with [γ-^32^P]-ATP (3000 Ci/mmol, Perkin Elmer) and T4 polynucleotide kinase (PNK; Thermo Fisher, #EK0031), according to standard procedures. Fluorescent substrates utilised 5′- and/or 3′-Cy5 or Cy3 labelled oligonucleotides. In the indicated cases, oligonucleotides containing three consecutive phosphorothioate (SP) linkages were used. The unpaired regions of these junctions are composed of heterologous sequences to prevent substrate dissociation by spontaneous branch migration. The strand composition of each substrate is detailed in Supplementary Table 2.

The plasmid DNA (pBluescript SK (-), pBSK, 2958 bp) was purified using commercial DNA purification kits (GenElute™ HP Plasmid Miniprep kit, Merck, #NA0160), according to standard procedures. To minimise plasmid nicking, which could affect subsequent D-loop formation that require negatively supercoiled plasmids^8,70^, the cell pellet was resuspended by pipetting and centrifugation steps were carried out at 8000 x *g*.

### Endonuclease assays with oligonucleotide-based substrates

For the experiments using radioactively labelled oligonucleotide-based D-loops (Fig. 1 and Supplementary Fig. 1), 10 nM protein was incubated with ∼1 nM 5’-^32^P end-labelled substrate in 25 µL of reaction buffer (Yen1 reaction buffer: 50 mM Tris-HCl pH 7.5, 0.5 mM MgCl_2_; Mus81-Mms4 reaction buffer: 25 mM Tris-HCl pH 7.5, 3 mM MgCl_2_, 100 mM NaCl, 0.1 mM DTT, 0.1 µg/mL BSA). In all reactions, enzymes represent 1/10 of the final volume (or enzyme storage buffer in mock reactions). After 10 min incubation at 30°C, 10 µL of each reaction was deproteinised by addition of 2 µL STOP solution (1.5% SDS, 10 mg/mL proteinase K (PK)) and incubation at 37°C for 1 h, followed by incorporation of 0.2 vol of 6x Ficoll loading buffer (15% Ficoll-400, 60 mM EDTA, 20 mM Tris-HCl pH 8.0, 0.5% SDS). Another aliquot of 10 µL was mixed with 1 vol of 2x denaturing loading buffer (1xTBE, 80% formamide) and incubated at 99°C for 3 min. The radiolabelled products were then separated by PAGE through 10% native or denaturing (7 M urea) PAGE gels in 1xTBE buffer. After electrophoresis, gels were dried onto 3MM Whatman chromatography paper (GE Healthcare), exposed to a phosphor screen (Fujifilm), visualised in a Typhoon FLA9500 (GE Healthcare), and quantified by densitometry using ImageQuant software (GE Healthcare).

For kinetics experiments using fluorescently labelled oligonucleotide-based D-loops (D3, D3-SP and nicked D3) (Fig. 2), 20 nM protein was incubated with 10 nM 5′-Cy5 and 5′-Cy3 labelled substrate. Reactions were performed as described above. Aliquots were withdrawn at the indicated times (0, 0.5, 1, 2, 4, 8, 16, 32, 64, 128 min) and analysed by 10% native and denaturing (7 M urea) PAGE in 1xTBE buffer. Fresh gels were scanned in a Typhoon FLA9500 and quantified by densitometry using ImageQuant software. Each experiment was done in triplicate.

For experiments using the long 3′-flap substrate, 2 µL of the indicated final concentration of RPA (25, 50, 100, 200, 400, and 600 nM) was pre-incubated with 40 nM 5′-Cy5-labelled 3′-flap for 10 min at 37°C in 8 µL Mus81-D-loop buffer (see next section). Then, 1 µL of Mus81-Mms4 at 400 nM final concentration was added and incubated for another 10 min at 30°C. Reactions were deproteinised by the addition of 2 µL STOP solution (1.5% SDS, 10 mg/mL proteinase K (PK)) and incubation at 37°C for 1 h, followed by incorporation of 0.2 vol 6x Ficoll loading buffer. Reaction products were analysed through 10% native PAGE in 1xTBE buffer. Fresh gels were scanned in a Typhoon FLA9500 and quantified by densitometry using ImageQuant software.

### Endonuclease assays with plasmid-based D-loops

#### Rad51/Rad54-mediated D-loops

To form the different D-loop structures, 40 nM of the indicated fluorescently labelled molecule was incubated with 2 µM Rad51 in D-loop buffer (35 mM Tris-HCl pH 7.5, 50 mM KCl, 1 mM DTT, 2 mM ATP, 20 mM creatine phosphate, 20 μg/mL creatine kinase), and 2.5 mM MgCl_2_ for Mus81-Mms4 reactions or 1.5 mM MgCl_2_ for Yen1 reactions) for 5 min at 37°C. When appropriate, RPA or SSB (600 nM) were added to the nucleoprotein filament and incubated for another 4 min, followed by the addition of Rad54 (300 nM) and incubation at 23°C for 3 min. D-loop formation was initiated by the addition of 2 µL pBSK (1/5 total volume, 64 ng/µL) in 10 μL final reaction volume and incubated for 5 min at 23°C. After D-loop formation, 1 µL at the indicated concentration of Yen1, Mus81-Mms4, the nuclease-dead mutants (always at the highest concentration used for the catalytically active enzyme), or storage buffer were added to the reaction followed by incubation at 30°C for the indicated times. Reactions were then deproteinised by the addition of 1.5 µL STOP solution and incubated at 37°C for 10 min, followed by the addition of 0.2 vol 6x glycerol loading buffer (66% Glycerol, 66 mM EDTA, 11 mM Tris-HCl pH 7.5). Reactions were analysed by electrophoresis in a 0.9% agarose gel in 1xTAE buffer developed at 90 V for 30 min. Fresh gels were imaged on a Typhoon FLA9500 and quantified by densitometry using ImageQuant software. Each experiment was done in triplicate.

When D1-3SP invading molecule was used, reactions were performed as described above, but increasing the final reaction volume to 25 µL. Then, 10 µL of each reaction was deproteinised and analysed on agarose gels. Another 10 µL was mixed with 1 vol of 2x denaturing loading buffer and incubated at 99°C for 3 min. Reaction products were then separated by 16% denaturing (7 M urea) PAGE gels in 1xTBE buffer. Gels were scanned on a Typhoon FLA9500 and quantified by densitometry using ImageQuant software.

#### RecA-mediated D-loops

RecA D-loops were generated in a similar way and in the same D-loop buffer as stated above, except in the presence of 15 mM MgCl_2_. D1 oligonucleotide (40 nM) was incubated with 4 µM RecA for 5 min at 37°C, followed by the addition of a single-stranded binding protein (100 mM SSB or RPA) when required. The reaction continued for an additional 5 min at 37°C. D-loop formation was initiated by the incorporation of pBSK (1/5 total volume, 64 ng/µL final concentration) and, after 0.5 min, reactions were diluted 1:10 in Yen1 buffer (50 mM Tris-HCl pH 7.5) for Yen1 reactions or 1:5 in Mus81 buffer (25 mM Tris-HCl pH 7.5, 100 mM NaCl) for Mus81-Mms4 reactions, to reach, in both cases, the optimal MgCl_2_ concentration for nuclease activity. One µL of each SSE (storage buffer for control reactions) was added at the indicated concentration and incubated for 5 min at 30°C. Reactions were then deproteinised by addition of 2.5 µL STOP solution (0.1 % SDS, 2 mg/mL PK) and incubation at 37°C for 2 h. Analysis was performed as described above. Each experiment was done three independent times.

#### Deproteinised D-loops

Rad51/Rad54-mediated D-loop formation was carried out as previously described. Before SSE incorporation, half of the reaction was deproteinised by the addition of 0.75 µL STOP solution and incubation at 37°C for 15 min. To remove PK and SDS, 1 vol of phenol-chloroform was added followed by DNA precipitation with 2 vol of 100% EtOH, 0.1 vol of 3 M sodium acetate pH 5.2, and 0.02 vol of 5 mg/mL glycogen. Pellet was resuspended in the original volume in which the D-loop reaction had taken place. Deproteinised D-loops were then used in nuclease reactions that were carried out and analysed as described above. After being scanned to visualise the fluorescent products, gels were stained with ethidium bromide (5 µL of 10 mg/mL ethidium bromide (EtBr) solution in 100 mL TAE) and imaged in a Gel Doc XR + System using the Image Lab software (Bio-Rad). Each experiment was carried out in triplicate.

#### D-loops with a stable presynaptic filament

For experiments using the non-hydrolysable ATP analogue ATPγS, Rad51/Rad54-mediated D-loops were generated as described above, with the following modifications: 40 nM D1 oligonucleotide was incubated with Rad51 (2 µM) in D-loop buffer without the ATP regeneration system (creatine phosphate and creatine kinase) and with 2.5 mM ATPγS or ATP for 9 min at 37°C, followed by addition of Rad54 (300 nM), 2.5 mM ATP, and incubation at 23°C for 3 min. D-loop formation was initiated by the incorporation of pBSK (640 ng), incubated for another 10 min at 23°C, and analysed as previously described. All experiments were carried out in triplicate.

### Mapping SSE cleavage sites on enzymatically generated D-loops

To map the cleavage sites in the invading molecule, Rad51/Rad54-mediated D-loop formation and treatment with the nucleases were done as described above, except in 25 µL total reaction volume. After nuclease incubation, 10 µL of each reaction were deproteinised and analysed on agarose gels as stated above, to monitor D-loop formation and nuclease activity. Another 10 µL aliquot was mixed with 1 vol of 2x denaturing loading buffer and incubated at 99°C for 3 min. Reaction products were then separated by denaturing PAGE (7 M urea) in 1xTBE buffer. Gels were scanned on a Typhoon FLA9500 and quantified by densitometry using ImageQuant software.

To analyse the incision in the donor plasmid molecule (pBSK), Rad51/Rad54-mediated D-loop formation (20 µL final volume) and cleavage with the indicated nuclease (as described), was followed by cleavage with 10 U *Bsp*HI (New England Biolabs, #R0517S) and incubated at 37°C for 20 min followed by dephosphorylation with 1 U Shrimp Alkaline Phosphatase (rSAP, New England Biolabs, #M0371S) and incubation for another 20 min. To verify D-loop formation and nuclease activity, half of each reaction was deproteinised and analysed on agarose gels as described above. The other half was denatured at 99°C for 3 min, followed by labelling using 1 µL [γ-^32^P]-ATP (3000 Ci/mmol) and 10 U T4 PNK for 1 h at 37°C. The non-incorporated isotope was removed using G-25 columns (GE Healthcare) and the eluted DNA was precipitated with 2 vol of EtOH, 0.1 vol of 3 M sodium acetate pH 5.2 and 0.02 vol of 5 mg/mL glycogen. The pellet was resuspended in 10 µL 2x denaturing loading buffer and samples were analysed through denaturing PAGE, exposed to a phosphor screen, and visualised in a Typhoon FLA9500.

For mapping experiments using Sanger sequencing, D-loop formation in 25 µL final volume and incubation with the indicated nuclease, each reaction was deproteinised by the addition of 3.75 µL STOP solution and incubation at 37°C for 30 min. To verify D-loop formation and nuclease activity, 5 µL of the reaction were analysed on agarose gels. The rest of the reaction was adjusted to 75 µL by water, followed by 1 vol of phenol-chloroform. After phenol-chloroform extraction, DNA precipitation was carried out with 2 vol of EtOH, 0.1 vol of 3 M sodium acetate pH 5.2, and 0.02 vol of 5 mg/mL glycogen. The pellet was resuspended in 22 µL 10 mM Tris-HCl pH 8.0. Finally, 10 µL was used for sequencing with the forward primer (SEQ-FW, from 1767 to 1784 bp) and the other 10 µL for sequencing with the reverse primer (SEQ-RV, from 2162 to 2177 bp) (Supplementary Table 1) in Stabvida laboratories.

### Detection of half-crossover precursors

In experiments detecting the formation of half-crossovers precursors, Rad51/Rad54-mediated D-loop formation was performed in 1.5 mM MgCl_2_ D-loop buffer, followed by simultaneous addition of 100 nM of both endonucleases and incubated for 1 h at 30°C. When appropriate, 5 U of T4 ligase (Thermo Fisher, #EL0011) was added and incubated for 1 h at 23°C. Then, 20 U of either *Eco*RV (EV, Thermo Fisher, #FD0304) or *Ahd*I (AI, New England Biolabs, #R0584S) was included and incubated at 37°C for 1 h. After treatment with the restriction enzymes, reactions were deproteinised, analysed by agarose gel, scanned in a Typhoon FLA9500, and stained with EtBr as stated above. For analysis by denaturing PAGE (using 7 M urea), after the cleavage with restriction enzymes, reactions were mixed with 1 vol of 2x denaturing loading buffer and incubated at 99°C for 3 min. Reaction products were then separated by denaturing PAGE in 1xTBE buffer and fresh gels were scanned on a Typhoon FLA9500.

### Statistical analysis

Unless indicated otherwise, data is presented as the mean of three independent experiments ± SD. Statistical analysis was performed using the student two-tailed T-test for samples with equal variance and the Welch T-test for samples with unequal variance. A p-value of less than 0.05 was considered statistically significant. *p < 0.05. **p<0.01. ***p<0.001. ****p<0.0001. n.s: not significant.

### Data availability

The data supporting the findings of this study are available within the article, its Supplementary Information files, and are available from the corresponding author upon request.

## Acknowledgements

The Blanco lab was supported from MCIN / AEI / 10.13039/501100011033 [PID2020-115472GB-I00]; MCIN / AEI / 10.13039/501100011033 / FEDER ‘Una manera de hacer Europa’ [BFU2016-78121-P]; Xunta de Galicia (XdG) / FEDER ‘Una manera de hacer Europa’ [ED431F-2016/019, ED431B-2016/016 and ED431C 2019/013]; CIMUS receives financial support from the XdG / FEDER [ED431G 2019/02, Centro Singular de Investigación de Galicia, accreditation 2019–2022]; F.J.A., R.C. and M.C. were recipients of pre-doctoral fellowships from XdG [ED481A-2015/011, ED481A-2018/041 and ED481A 2022/155]; M.C. was recipient of an undergraduate fellowship from AECC. The Krejci lab was supported from Czech Science Foundation (GACR 21-22593X); Masaryk University [MUNI/G/1594/2019]; European Union’s Horizon 2020 research and innovation programme under grant agreement No 812829; and European Structural and Investment Funds, Operational Programme Research, Development and Education “Preclinical Progression of New Organic Compounds with Targeted Biological Activity” (Preclinprogress, CZ.02.1.01/0.0/0.0/16_025/0007381).

## Author contributions

R.C., L.K. and M.G.B. designed the experiments. R.C. conducted most of the experimental part. F.J.A. performed preliminary mapping experiments on synthetic D-loops. M.C. performed some of the RecA experiments. M.S. purified some of the batches of yeast Rad51, RPA and Rad54. R.C. and M.G.B. wrote the manuscript. All authors read and approved the final manuscript.

## Additional information

**Supplementary information**

**Competing interest:** The authors declare no competing interest.

**Correspondence** and request for materials should be addressed to Lumir Krejci or Miguel G. Blanco

**Peer review information**

**Reprints and permission information**

## Supplementary Tables

**Supplementary Table 1.**
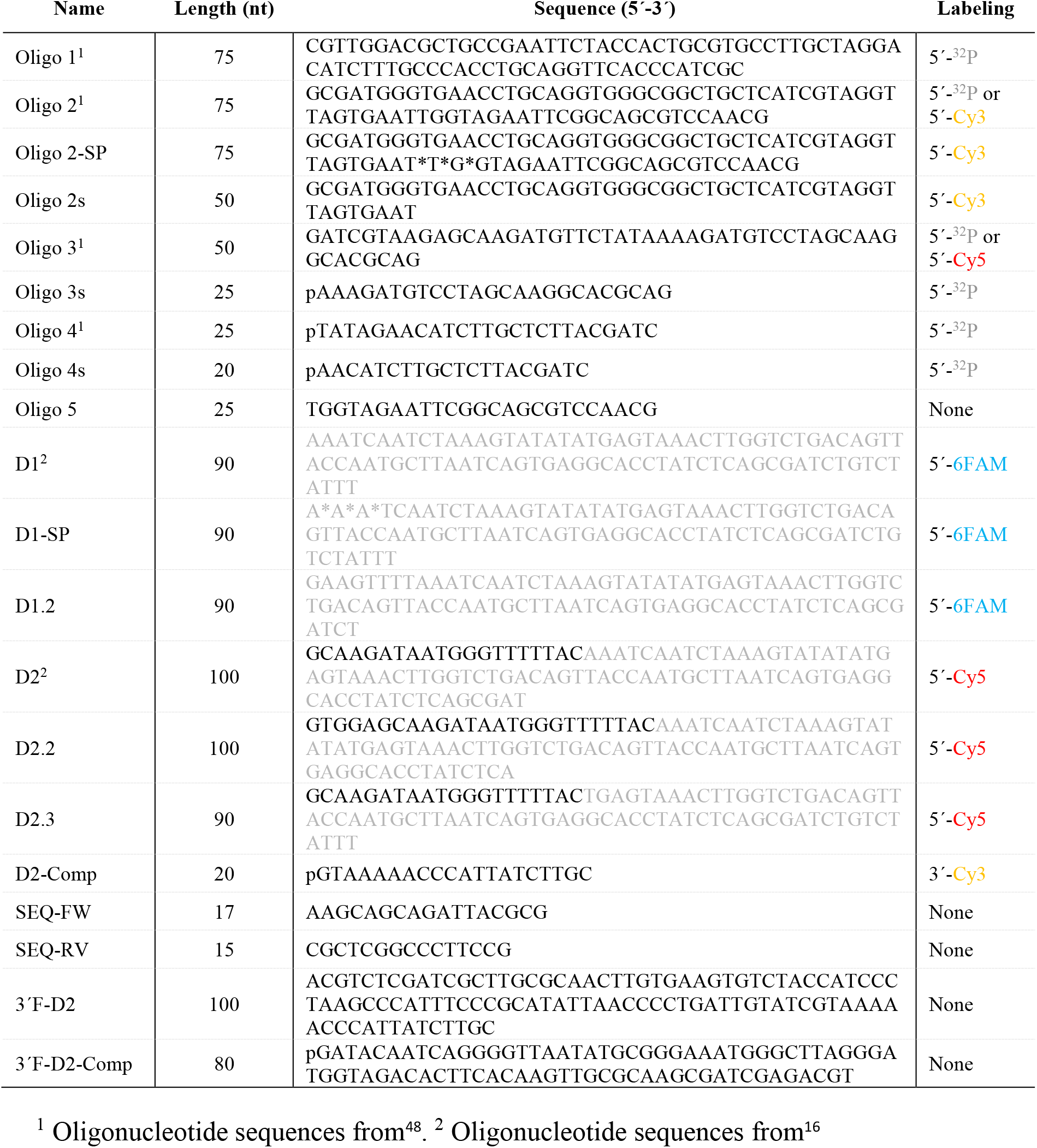
Oligonucleotide sequences used in this study. Labelling is specified when appropriate. The asterisk (*) indicates a phosphorothioate (SP) linkage. The presence of a phosphate group on the corresponding 5′-end of the oligonucleotide is denoted by a (p). Grey sequences indicate homology to pBSK plasmid.

**Supplementary Table 2.**
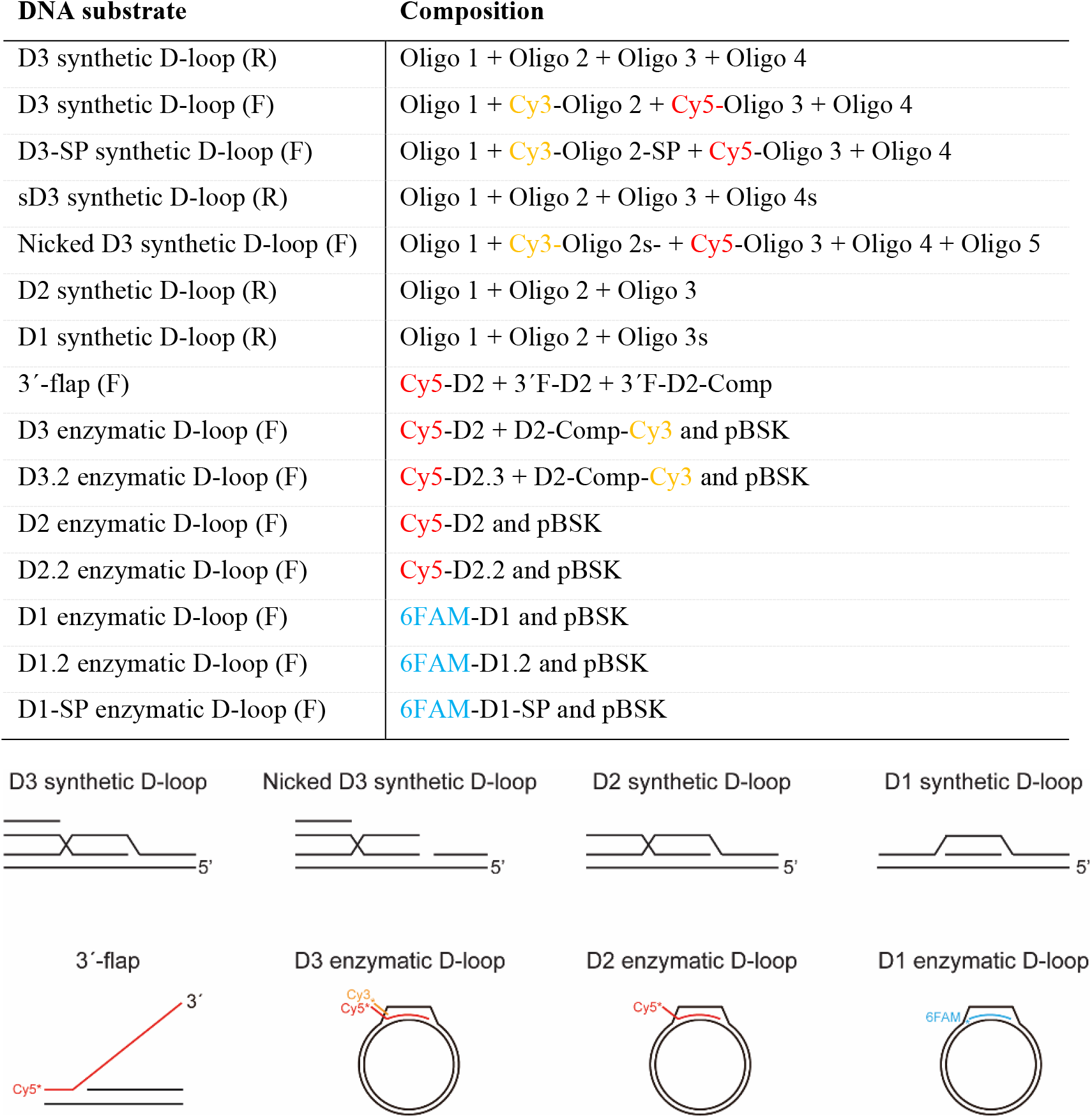
DNA substrate composition employing the oligonucleotides listed in Table 1. (R) denotes radioactive substrates, (F) denotes fluorescent substrates.

## Supplementary Figure legends

**Supplementary Figure 1.**
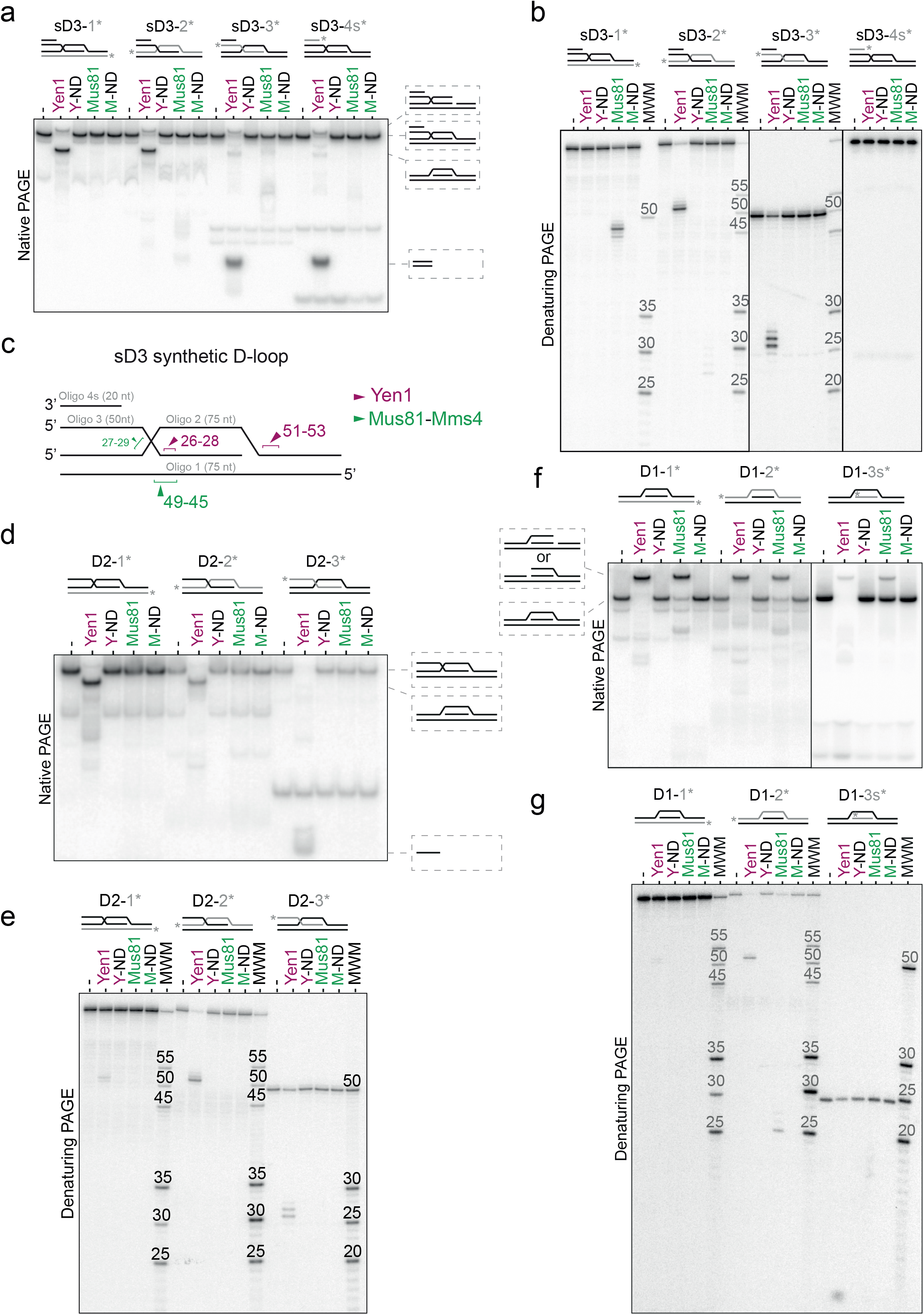
Exonuclease activity of Yen1 depends on a free-5′end at the branched point. (**a**) Synthetic shorter D3 D-loop (sD3) was labelled with ^32^P at the 5’-end (asterisk) of each strand (grey) and incubated with 10 nM Yen1^WT^ (Yen1), Yen1^ND^ (Y-ND), Mus81^WT^ (Mus81), or Mus81^ND^ (M-ND) for 10 min at 30°C. (-) indicates no enzyme. The reaction products were analysed using 10% native PAGE and scanned. Schematic representation of the substrate and cleavage products are depicted on the right. (**b**) The same reaction products as in (**a)** were analysed on 10% denaturing PAGE. A molecular weight marker (MWM) consisting of a mixture of 5’-^32^P end-labelled oligos of defined length was used (**c**) Schematic representation of Yen1 (purple) and Mus81 (green) incision sites on the short D3 substrate (sD3). (**d**) and (**e**) Reactions with D2 synthetic D-loop (D2) were carried out and labelled as in (**a)** and (**b)**. (**f**) and (**g**) Reactions with D1 synthetic D-loop (D1) were carried out and labelled as in (**a)** and (**b**).

**Supplementary Figure 2.**
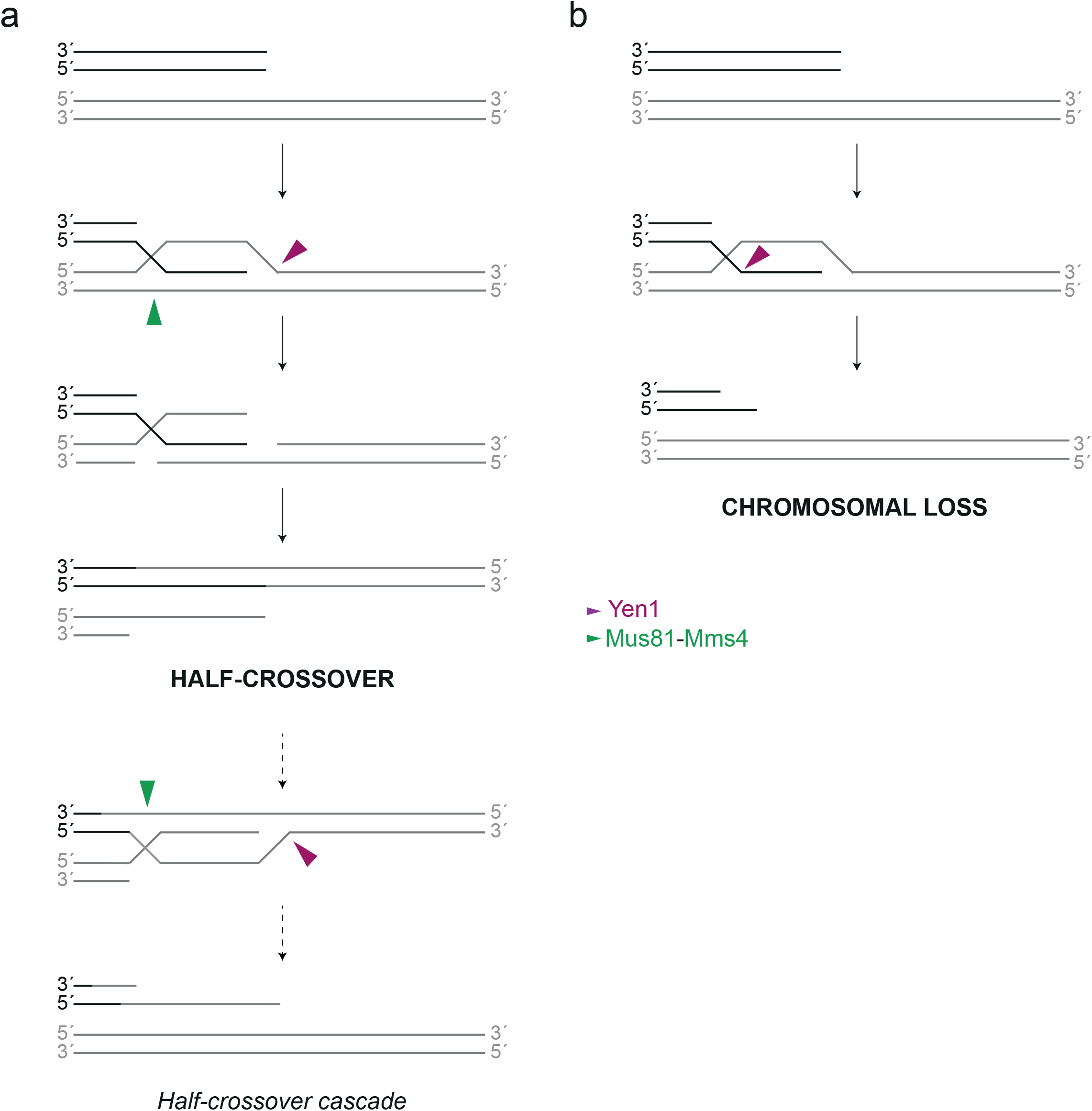
Graphical representation of the generation of half-crossovers and chromosomal loss events in the context of one-ended DSB repair by BIR. (**a**) Generation of a half-crossover: Mus81-Mms4 cleavage at the strand complementary to the invading molecule, in combination with Yen1 incision at the end of the displaced strand on a nascent D-loop, can lead to the formation of a half-crossover. Note that Mus81-Mms4 incision requires DNA synthesis for ligation to occur. This event results in a formation of a new one-ended DSB that has the potential to initiate further rounds of D-loop formation and cleavage, leading to half-crossover cascades. Mus81-Mms4 incision is indicated by a green arrowhead, while Yen1 incision is indicated by a purple arrowhead. (**b**) Chromosomal loss: Yen1 incisions at the invading strand can result in a chromosomal loss event, where a portion of the chromosome is lost due to degradation of the cleaved DNA fragment.

**Supplementary Figure 3.**
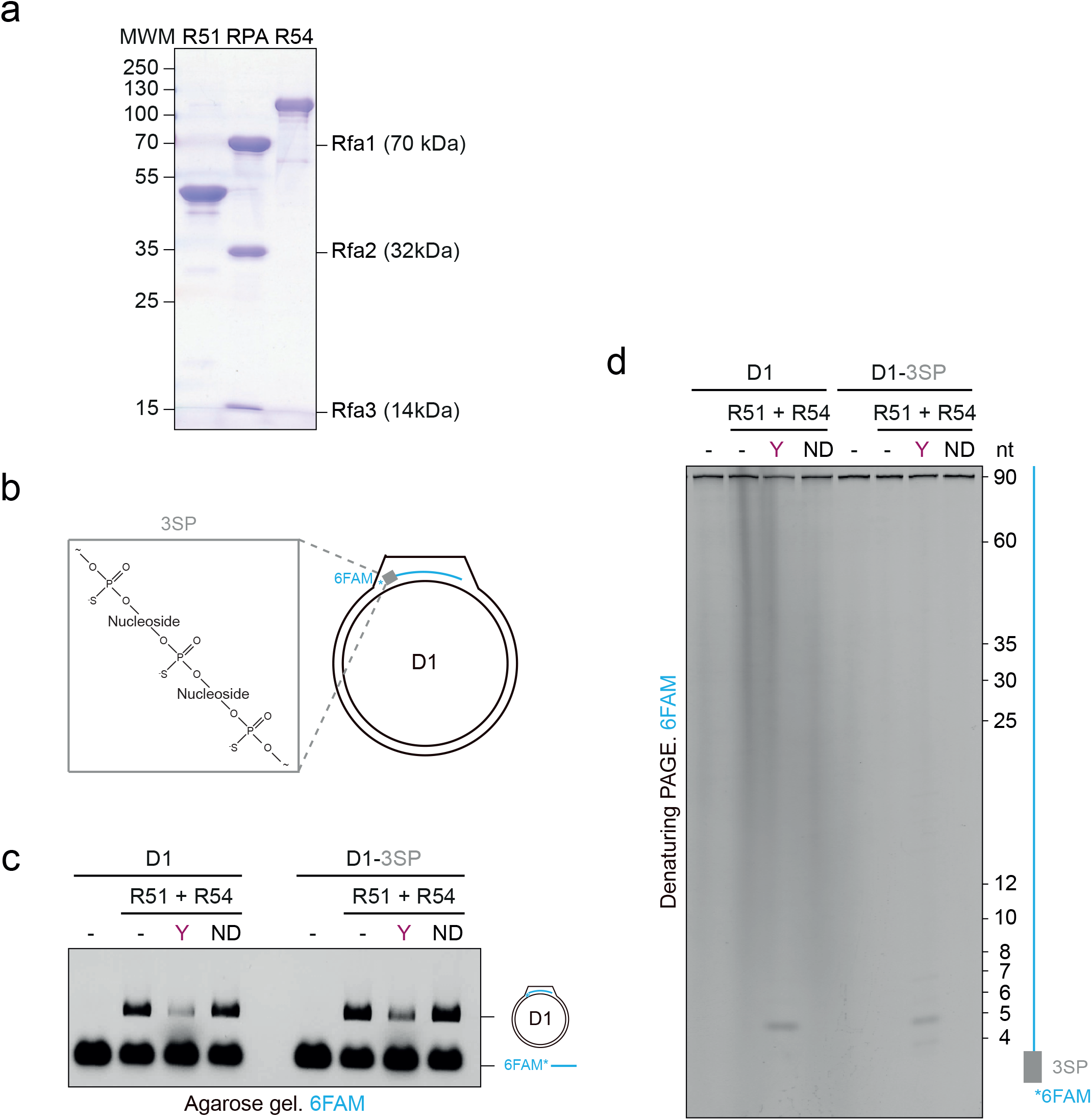
Yen1 can process the 5′end of the fully homologous invading strand. (**a**) Yeast Rad51, RPA, and Rad54 were purified, analysed by SDS-PAGE, and stained with Coomassie. Molecular weight markers are indicated in kDa. (**b**) Schematic representation of a D1 D-loop containing a D1 oligo with 3 phosphorothioate (SP) linkages at the 5′-end (D1-3SP, SP linkages between nt 1-2-3-4, indicated with a grey box). (**c**) Analysis of Yen1 cleavage of D1 and D1-3SP D-loops. Reactions were carried out as described in Figure 3 using 400 nM Yen1 (Y) or Yen1^ND^ (ND). (-) indicates no enzyme; R51, Rad51; R54, Rad54. Reaction products were analysed by native agarose gel electrophoresis and scanned. (**d**) Same as (**b**), but cleavage products were analysed on a 16% denaturing PAGE. A mixture of 5’-6FAM-end-labelled oligos of defined length was used as marker.

**Supplementary Figure 4.**
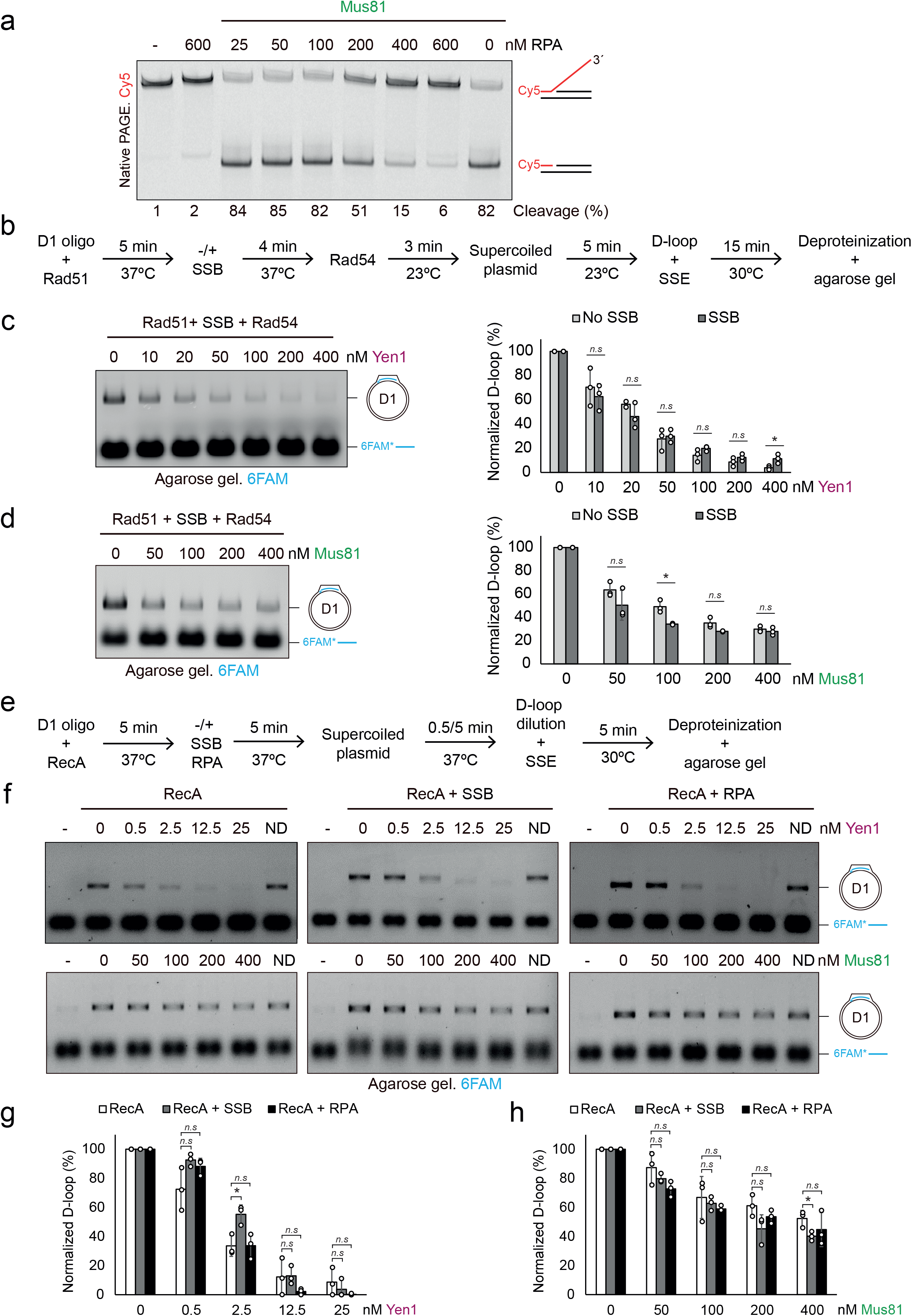
Yen1 and Mus81 cleavage in the presence of *E.coli* SSB or RecA proteins. (**a**) Effect of RPA on Mus81 cleavage using a 3′-flap. 3′-flap DNA (40 nM) was preincubated with the indicated RPA concentrations for 10 min at 37°C. Then Mus81 (400 nM) was added to the reaction and incubated for 10 min at 30°C. Reactions were deproteinised, analysed on a 10% native PAGE, and scanned. (-) indicates no enzyme. Cleavage products are depicted on the right. (**b**) Scheme of Rad51/Rad54-mediated D1 D-loop reactions with SSB protein. Reactions were carried out as in Fig. 4a with 600 nM SSB. (**c**) Representative agarose gel from reactions with Yen1 and enzymatic D1 D-loop with SSB. Reactions were carried out, visualised, and plotted as in Fig. 4b. (**d**) Same as (**c**) but using Mus81. (**e**) Scheme of RecA-mediated D1 D-loop reactions with SSB or RPA. 5′ 6FAM-end-labelled D1 oligonucleotide (40 nM) was incubated with RecA (4 µM) at 37°C for 5 min. When appropriate, SSB or RPA (100 nM) was added and incubated for 5 min at 37°C. Then, supercoiled pBSK (640 ng) was added and incubated at 37°C for 0.5 min in reactions with RecA and 5 min in reactions with RecA + SSB or RPA. D-loop reactions were diluted 1:10 prior to Yen1 incorporation or 1:5 prior to Mus81 incorporation. The indicated SSE concentration was added, and reactions were incubated at 30°C for 5 min. (**f**) Representative agarose gels from reactions with Yen1 (top) or Mus81 (bottom), and RecA-mediated D1 D-loop alone (left), with SSB (middle) or with RPA (right). (-) indicates no enzyme. (**g**) Quantification of D-loop cleavage by Yen1 is shown by normalizing D-loops from (**f**), setting the initial D-loop yield as 100%. Plotted are means ± SD (n = 3). White circles represent individual values. (**h**) Same as (**g**), but using Mus81.

**Supplementary Figure 5.**
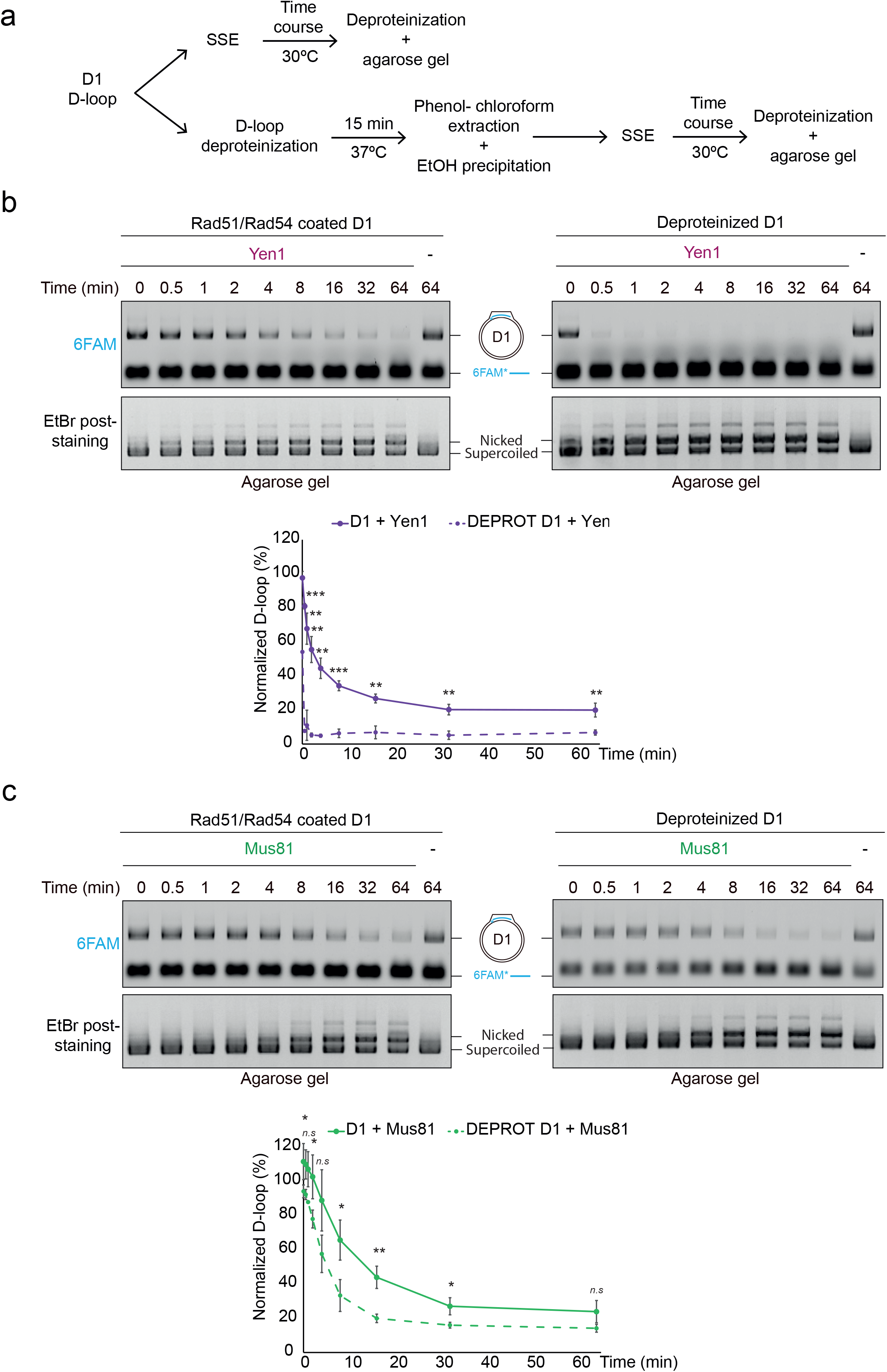
Effect of D-loop deproteinization on nuclease cleavage. (**a**) Experimental scheme of nuclease reactions on Rad51/Rad54-coated and deproteinised D1 D-loops. After Rad51/Rad54-mediated D1 D-loop formation, half of the reaction was directly incubated with 400 nM SSE. The other half was deproteinised for 15 min at 37°C, followed by DNA purification using phenol-chloroform extraction and EtOH precipitation. Subsequently, SSE (400 nM) was added and incubated at 30°C. Aliquots were removed at the indicated times and analysed on agarose gel. (**b**) Representative agarose gels of Yen1 cleavage on a Rad51/Rad54-coated (left) or deproteinised (right) D1 D-loop. Gels were scanned by Typhoon FLA9500 (top), followed by post-staining with EtBr (bottom). (-) indicates no nuclease. Quantification of D-loop cleavage by Yen1 is shown at the bottom. D-loops were normalised by setting the initial D-loop yield as 100%. Plotted are means ± SD (n = 3). Solid line: Rad51/Rad54-coated D-loops. Dashed line: deproteinised D-loops. (**c**) Same as (**b**), but for 400 nM Mus81.

**Supplementary Figure 6.**
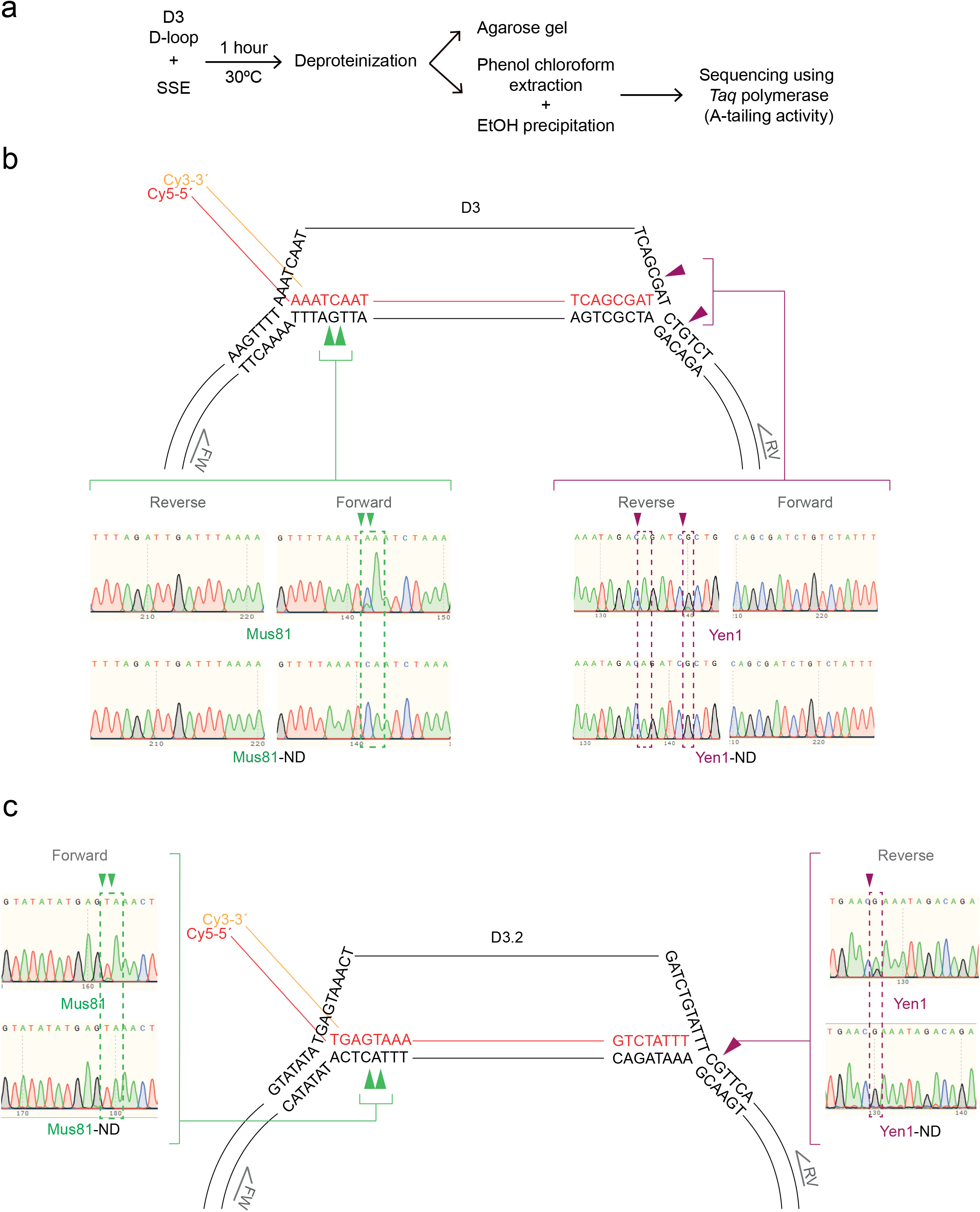
Mapping of cleavage site of Yen1 and Mus81 in Rad51-mediated D3 D-loops using sequencing. (**a**) Scheme of the mapping strategy using DNA sequencing. After D-loop formation, 400 nM Yen1 or Mus81 was added and incubated for 1 h at 30°C, followed by deproteinisation. The DNA was purified by phenol-chloroform extraction, followed by EtOH precipitation. Reaction products were sequenced with *Taq* polymerase using forward and reverse primers. (**b**) Graphical representation of the incisions mapped on plasmid-based D3 D-loops. The nucleotide sequence near the branch point is shown. Chromatograms derived from sequencing using forward and reverse primers are depicted at the bottom. Left: reactions with Mus81. Green arrows indicate Mus81 cleavage sites. Adenine incorporation after the cleavage point is indicated with a green dotted box. Nuclease-dead mutant controls are shown (ND). Right: Reactions with Yen1. Purple arrows indicate Yen1 cleavage sites and adenine incorporation is indicated with a purple dotted box. Nuclease-dead mutant controls are shown (ND). (**c**) Same as (**b**) but using D3.2 oligo for D-loop formation. Note that both branched points are different from the previous ones (from position 1952 to 2022 of pBSK). Only chromatograms of the region of interest are shown.

**Supplementary Figure 7.**
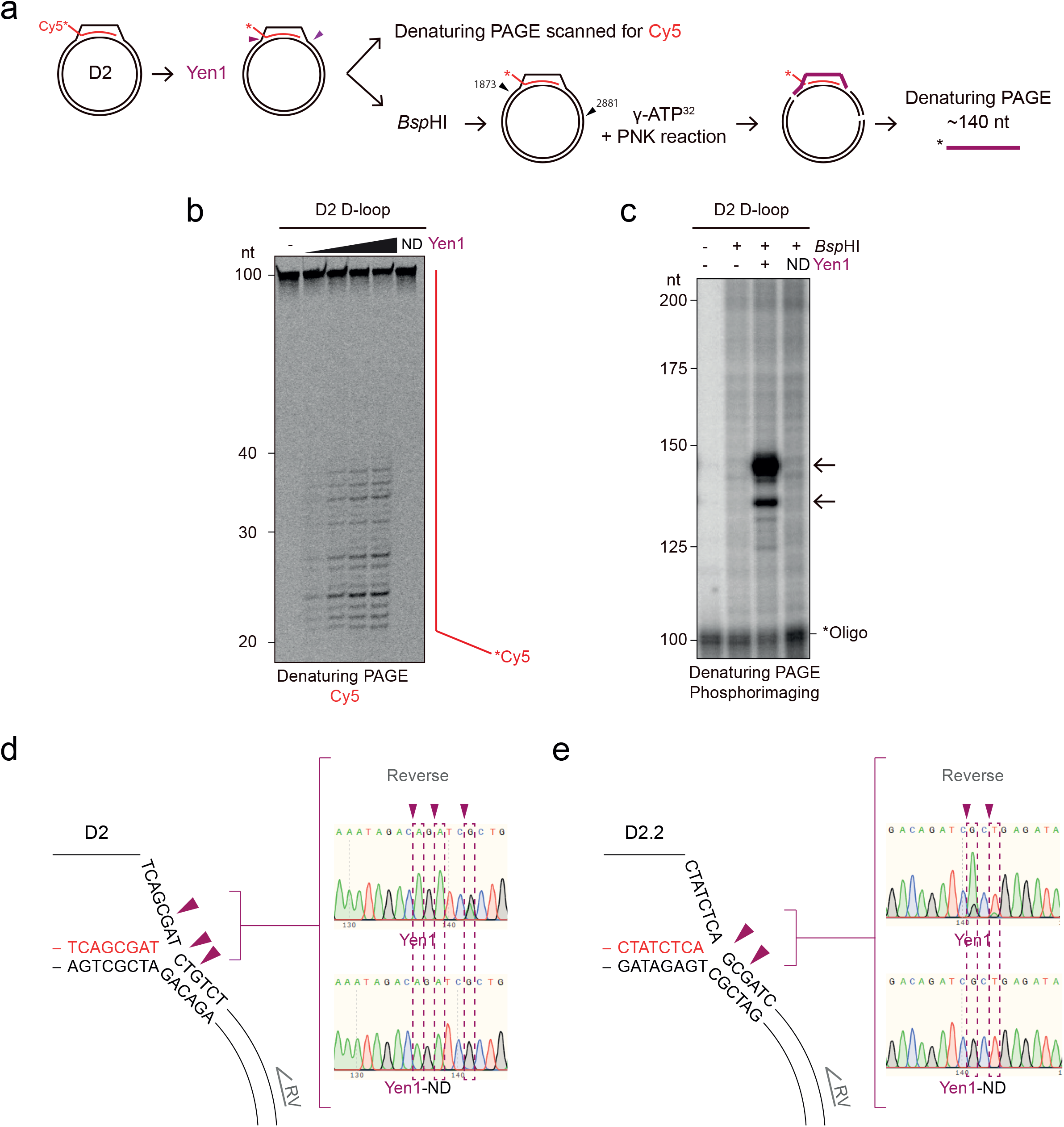
Mapping of Yen1 cleavage sites within D2 D-loops. (**a**) Mapping strategy for the D2 D-loop structure. The invading oligo is homologous to positions 1932 to 2012 of pBSK. After D2 D-loop formation, SSE activity on the invading molecule was analysed by fluorescence scanning after denaturing PAGE; SSE activity on the plasmid was analysed after *Bsp*HI treatment, radioactive labelling, and analysis on denaturing PAGE by phosphorimaging. (**b**) To identify the cleavage site in the invading molecule after D2 D-loop formation, increasing concentrations of Yen1 (50, 100, 200, 400 nM) or 400 nM Yen1^ND^ (ND) were added to the reaction and incubated for 1 h at 30°C. (-) indicates no enzyme. Reaction products were analysed by 12% denaturing PAGE and scanned. A mixture of 5’-6FAM-end-labelled oligos of defined length was used as marker. (**c**) To identify incisions within the plasmid, after D2 D-loop formation, 400 nM Yen1 was added to the reaction and incubated for 1 h at 30°C. Then, pBSK was digested with 10 U *Bsp*HI at 37°C for 20 min, followed by treatment with rSAP for another 20 min at 37°C and heat inactivation at 99°C for 5 min. Reaction products were labelled using T4 PNK and ^32^P-γ-ATP and analysed by 6% denaturing PAGE followed by phosphorimaging. A mixture of DNA fragments was radioactively labelled and used as marker. Arrows indicate products of expected sizes. (**d**) Graphical representation of the incisions mapped on plasmid-based D2 D-loops. The nucleotide sequence near the branch point is shown. Sequencing reactions were carried out as in Supplementary Fig. 5. Chromatograms derived from sequencing using the reverse primer are depicted on the right. Note that only chromatograms of the region of interest are shown. Purple arrows indicate Yen1 cleavage sites and adenine incorporation is pointed out with a purple dotted box. Nuclease-dead mutant controls are shown (ND). (**e**) Same as (**d**) but employing a D2.2 oligo for D-loop formation. Note that the second branched point is different from the previous one (from position 2012 to 2007 of pBKS).

**Supplementary Figure 8.**
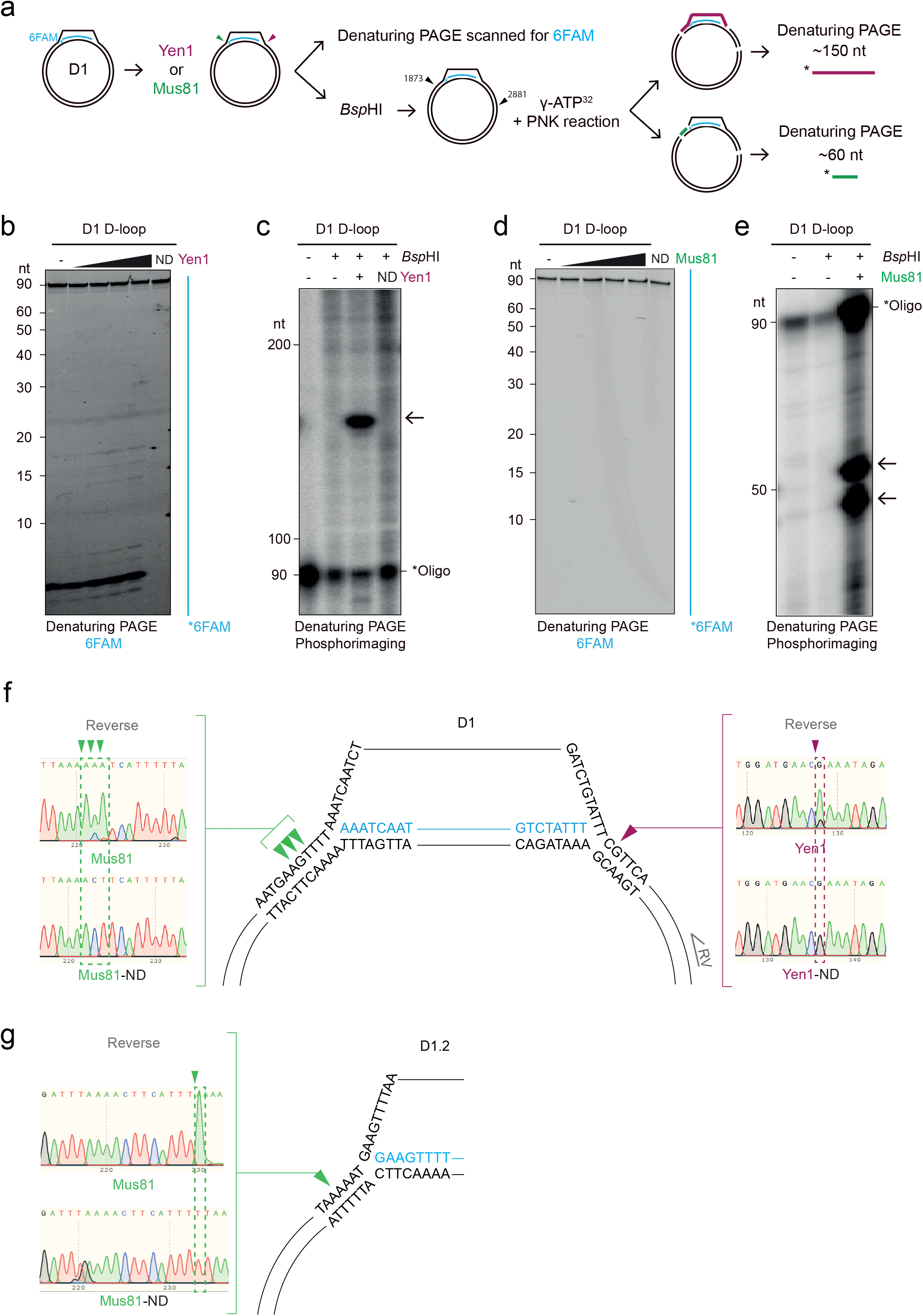
Mapping Yen1 and Mus81 the cleavage sites within D1 D-loops. (**a**) Mapping strategy for the D1 D-loop structure. The invading oligo is homologous to positions 1932 to 2022 of pBSK. After D1 D-loop formation, SSE activity on the invading molecule was analysed by fluorescence scanning after denaturing PAGE; SSE activity on the plasmid was analysed after *Bsp*HI treatment, radioactive labelling, and analysis on denaturing PAGE by phosphorimaging. (**b**) To identify cuts in the invading molecule, after D1 D-loop formation, increasing concentrations of Yen1 (50, 100, 200, 400 nM) or 400 nM Yen1^ND^ (ND) were added to the reaction and incubated for 1 h at 30°C. (-) indicates no enzyme. Reaction products were analysed by 16% denaturing PAGE and scanned a. A mixture of 5’-6FAM-end-labelled oligos of defined length was used as marker. (**c**) To identify incisions within the plasmid, after D1 D-loop formation, 400 nM Yen1 was added to the reaction and incubated for 1 h at 30°C. Then, pBSK was digested with 10 U *Bsp*HI at 37°C for 20 min, followed by treatment with rSAP for another 20 min at 37°C and heat inactivation at 99°C for 5 min. Reaction products were labelled using T4 PNK and ^32^P-γ-ATP and analysed by 6% denaturing PAGE followed by phosphorimaging. A mixture of DNA fragments was radioactively labelled and used as marker. The arrow indicates product of expected size. (**d**) and (**e**) Same as (**b**) and (**c**), but using Mus81. (**f**) Graphical representation of the incisions mapped on plasmid-based D1 D-loops. The nucleotide sequence near the branch point is shown. Sequencing reactions were carried out as in Supplementary Fig. 5. Chromatograms derived from sequencing using the reverse primers are shown. Note that only chromatograms of the region of interest are shown. Left: reactions with Mus81. Green arrows indicate Mus81 cleavage sites. Adenine incorporation after the cleavage point is indicated with a green dotted box. Nuclease-dead mutant controls are shown (ND). Right: Reactions with Yen1. Purple arrows indicate Yen1 cleavage sites and adenine incorporation is pointed out with a purple dotted box. Nuclease-dead mutant controls are shown (ND). (**g**) Same as (**f**), but employing a derivative D1.2 invading oligo. Note that the first branched point is different from the previous one (from position 1932 to 1924 nt of pBSK).

